# Leg compliance is required to explain the ground reaction force patterns and speed ranges in different gaits

**DOI:** 10.1101/2024.09.23.612940

**Authors:** Ali Tehrani Safa, Tirthabir Biswas, Arun Ramakrishnan, Vikas Bhandawat

**Affiliations:** School of Biomedical Engineering, Science and Health Systems Drexel University Philadelphia, PA 19104, USA; Department of Neurobiology Northwestern University Evanston, IL 60208, USA; Janelia Research Campus Howard Hughes Medical Institute Ashburn, VA 20147, USA; College of Nursing and Health Professionals Drexel University Philadelphia, PA 19104, USA

## Abstract

Two simple models – vaulting over stiff legs and rebounding over compliant legs – are employed to describe the mechanics of legged locomotion. It is agreed that compliant legs are necessary for describing running and that legs are compliant while walking. Despite this agreement, stiff legs continue to be employed to model walking under the assumption that the compliance of the leg during walking is high enough to be considered stiff. Here we study gait choice and walk-to-run transition in a biped with compliance and show that the principles underlying gait choice and transition are completely different from stiff legs. Two findings underpin our conclusions: First, at the same speed, step length, and stance duration, multiple gaits that differ in the number of leg contraction cycles are possible. Among them, humans and other animals choose the (normal) gait with M-shaped vertical ground reaction forces (vGRF) not just because of energy considerations but also constraints from forces. Second, the transition from walking to running occurs because of three factors: vGRF minimum at mid-stance characteristic of normal walking, synchronization of horizontal and vertical motions during single support, and velocity redirection during the double support. The insight above required an analytical approximation of the double spring-loaded pendulum (DSLIP) model describing the intricate oscillatory dynamics that relate single and double support phases. Additionally, we also examined DSLIP as a quantitative model for locomotion and conclude that DSLIP speed-range is limited. However, insights gleaned from the analytical treatment of DSLIP are general and will inform the construction of more accurate models of walking.

## 1 Introduction

The CoM movement and the forces exerted on them follow relatively simple patterns conserved across animals, suggesting that the overall animal-substrate interactions and, therefore, the underlying mechanical principles are simple and general (Blickhan, 1989; Geyer et al., 2006). The best example of this generality is observed during running: Irrespective of the size of the animal, and the number of legs it uses, during running, the CoM reaches its minimum height at mid-stance and the vertical ground reaction force (vGRF) has an inverted “U”-shaped profile with a midstance maximum. This profile is well-explained by the spring-loaded inverted pendulum (SLIP), in which the mass of the animal is concentrated at a point and supported by a massless spring (Blickhan, 1989; McMahon and Cheng, 1990; Blickhan and Full, 1993; Ahn et al., 2004; Daley et al., 2007; Nishikawa et al., 2007; Schmitt, 1999).

Unlike running, it is unclear whether leg compliance is important for walking – the gait used at low speeds. Initially, the inverted pendulum (IP) model, which uses a non-compliant or rigid leg, was used to model walking (Griffin et al., 2004; Usherwood, 2005; Buczek et al., 2006). The IP model successfully models the energetics of walking (Kuo, 2001; Donelan et al., 2002; Kuo, 2002; Kuo et al., 2005) explaining correctly the exchange of kinetic and potential energy during walking: During the first half of the stance phase, the speed of the CoM decreases and height increases; the increased potential energy is reconverted into kinetic energy during the second half of the stance phase.

With modifications, IP can also model the work done during step-to-step transition. During walking, the CoM velocity vector is directed downward at the end of the step and must be redirected upward before the next step (Kuo, 2001; Adamczyk and Kuo, 2009; Donelan et al., 2002). In the IP model, velocity redirection occurs instantaneously, therefore, the work performed during the transition cannot be estimated. Regardless, many trends in the work performed during walking can be explained by distributing the force impulse in IP over a finite period of time; however, these modifications are entirely *ad hoc*. Another, perhaps more fundamental limitation of the IP model is that it cannot model the double-humped or M-shaped vertical GRF (vGRF) during walking.

This limitation has been addressed in many ways: by modeling non-impulsive impact forces at the beginning and end of each step, and by using a telescoping actuator with bounds on impact forces (Srinivasan and Ruina, 2006; Srinivasan, 2011). However, the model that produces the most naturalistic force profiles assumes a linear relation between force and leg length, implying that a linear spring is likely necessary to model vGRF during walking. Even with these modifications, IP model cannot model the mid-stance minimum in vGRF. This limitation - as shown in this study – is a significant flaw when considering walk-run transition. *It turns out that the mid-stance minimum, while making M-shaped walking an economical gait, is also a reason why the range of walking speeds is limited, and this is a key new insight we provide*.

The limitation of the IP model and the recent realization that legs are compliant during walking (Lee and Farley, 1998; Buczek et al., 2006) led to the development of the double SLIP (DSLIP) model, in which each leg of a biped is modeled as a spring (Figure 1A). DSLIP extends SLIP with a double stance phase during which CoM is supported by two ’springy’ legs (Geyer et al., 2006; Rummel et al., 2010). DSLIP can produce M-shaped GRFs observed during human walking by providing smooth velocity redirection during the double-stance phase. It also produces trajectories with mid-stance heights that are lower than IP and more in accordance with experimental data. Although DSLIP is an attractive model, there are several issues regarding DSLIP as a model. The first issue is whether DSLIP can explain the choice of gait at a given speed. Although it is clear that DSLIP is versatile and all the major gaits observed during bipedal walking can emerge from the DSLIP model (Gan et al., 2018), the speed range over which DSLIP supports M-shaped walking is limited compared to the range observed in animals (Geyer et al., 2006; Lipfert et al., 2012). The reasons for this limited range of speed supported by DSLIP are not understood. It is unclear whether this disparity in speed ranges is a fundamental limitation of the DSLIP model or a matter of quantitative detail.

**Figure 1.**
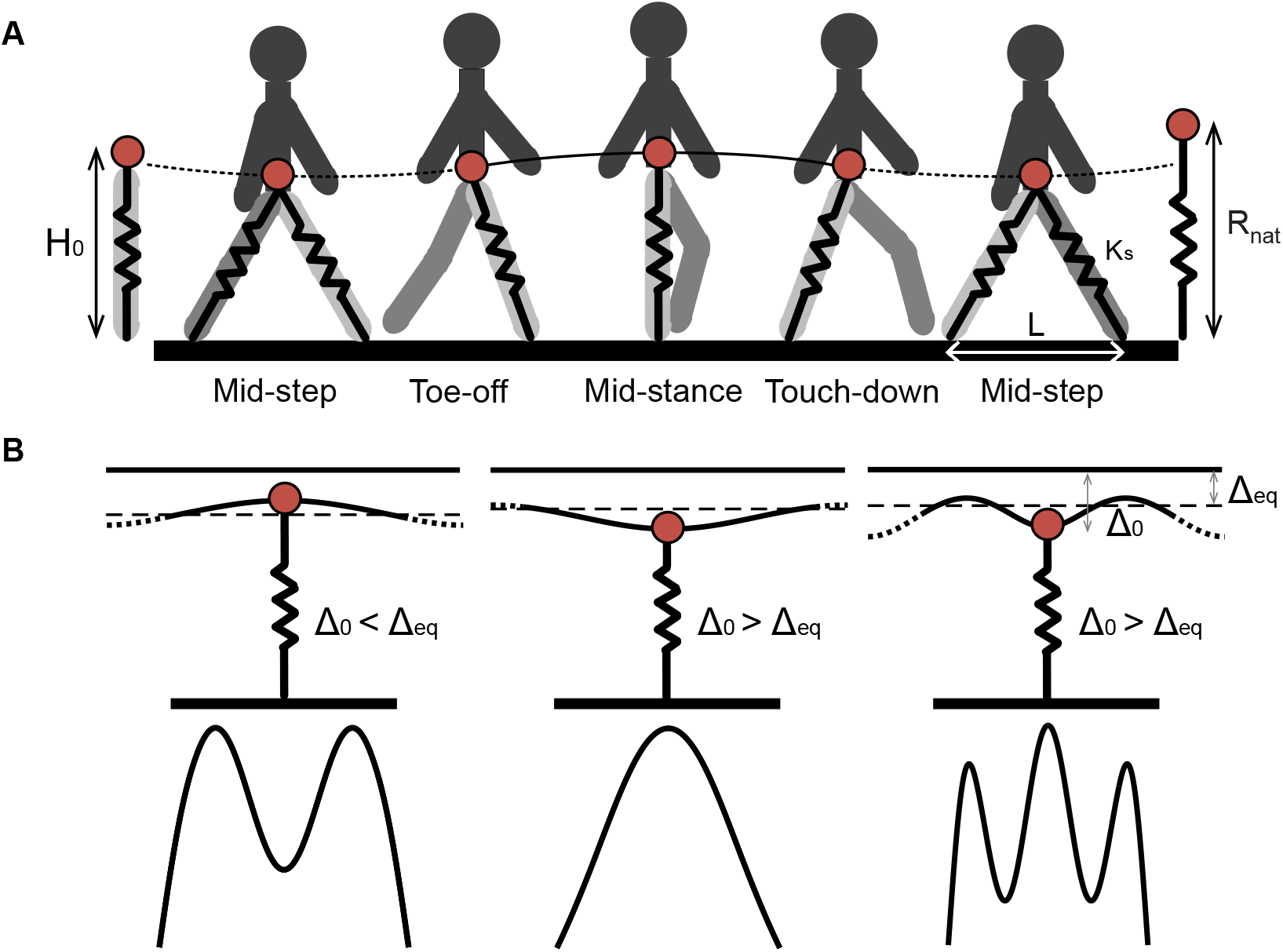
The position of the CoM at mid-stance in relation to the equilibrium point of the spring-mass system determines the GRF profile. **A**. Gait cycle for normal human walking showing the mid-stance maximum in height and a mid-step minimum in height. The DSLIP model is overlayed on the step cycle. Solid lines and dotted lines represent single and double stance phases respectively. **B**. The top row shows the position of the CoM at mid-stance in relation to the equilibrium position. During human walking (left), the CoM at mid-stance is above this equilibrium point; the resulting vGRF will be at its minimum and produce an M-shaped vGRF. In grounded running (middle), the CoM is at its minimum height below the equilibrium point resulting in the maximum spring contraction/force at mid-stance. Walking with multiple oscillations (right) can have either a maximum or minimum CoM height regarding the number of oscillations. Again with the same logic, the extremums of vGRF profile are defined based on the position of the CoM related to the equilibrium point.

A second related issue is that of gait transition from M-shaped walking. The issue of what mechanical principles underlie gait transitions in a DSLIP model or in a system with compliant legs has not been studied well enough (Srinivasan, 2011) and it is assumed that the reasons for gait transitions in DSLIP are the same as that in the IP model because the effective spring during walking is stiff. This assumption seems reasonable at face value, but has never been rigorously examined. In this study, we will show that the mechanical principles underlying the gait transition are completely different when compliance is added to the leg.

A third issue is how well DSLIP models the kinematics and mechanics of human walking. This question has not been evaluated rigorously. In studies in which DSLIP is compared to experimental data, it predicts within-step variations in CoM height and ground reaction forces (GRFs) (Lipfert et al., 2012; Hubel and Usherwood, 2015) that are larger than those observed experimentally. A larger issue is how a successful model is defined. Most studies focus on a single aspect of locomotion such as GRFs. Considering GRFs, CoM kinematics, and real non-dimensionalized time (and not normalized time) at the same time is crucial because the CoM height, *H*, along with the gravitational acceleration constant, *g* ≈ 9.8 m/s^2^, determines the natural timescale of the system 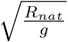,where, *R*_*nat*_ ≈ *H*, is the natural spring length. A successful model must produce realistic GRFs within the constraints of experimentally observed CoM kinematics and stance duration. These three constraints are rarely satisfied (Maus et al., 2010; Lipfert et al., 2012; Maus et al., 2015) simultaneously in most studies of locomotion, leaving the problem under-constrained. A previous study used this approach to model the single support phase of human walking (Antoniak et al., 2019).

These issues raise the question of whether adding compliance to the leg is worth the added complexity. In this study, we show that adding leg compliance through the DSLIP model is worth the complexity. During locomotion, the radial and angular motions of the CoM must be synchronized. Leg compliance provides a natural mechanistic basis for understanding the implications of this synchronization. We show that leg compliance explains the gaits observed at a given speed and how they relate to different oscillatory modes of the spring. *We further argue that the normal gait with the characteristic “M”-shaped GRF is preferred over other available walking gaits with different number of oscillatory cycles because of a combination of energy and force constraints. The other gaits either require more energy or unusually large forces, and this is another key insight of our work*. Surprisingly, the preference for the M-shaped vGRF also limits the range of speed over which walking is possible by requiring a large redirection of the velocity vector while simultaneously making it difficult to extract the required vertical forces necessary for the same. Thus, the reasons why a walker with compliant legs undergoes gait transition is fundamentally different from why a stiff-legged walker transitions. While the DSLIP model is particularly limited in its ability (likely because springs are assumed to be linear) to produce M-shaped GRFs, it does explain the fundamental reason why humans (and other animals) transition to faster gaits at size-specific speeds (Froude number ≡ (speed)^2^/*gR*_*nat*_) that are significantly lower than 1, the approximate transition speed predicted by IP.

## 2 Results

Throughout the manuscript, we employ the DSLIP model (Figure 1A) in which both legs are modeled as massless springs with the same stiffness, *K*_*s*_, and natural length, *R*_nat_. The dynamics of the single stance phase are the same as SLIP; the swing dynamics are not modeled. The single stance phase transitions to a double stance phase when the distance between the CoM and the future footstep equals the spring’s natural length, and we assume that the swing leg has “touched down”. We will focus on symmetric gaits so that the lift-off of the receding leg and the touch-down of the leading leg occur at time points given by time-reversal symmetry about the mid-step time. All variables in their dimensional and dimensionless forms are enumerated in the table below.

The model’s parameters

| Parameter | Symbol | Dimensionless form | Relation |
| --- | --- | --- | --- |
| Mass | M | N/A | N/A |
| Acceleration of gravity | g | N/A | N/A |
| Weight | W | N/A | $W = Mg$ |
| Natural spring length | $R_{nat}$ | N/A | N/A |
| Time | $t$ | $\tau$ | $\tau = t\sqrt{\frac{g}{R_{nat}}}$ |
| Single stance time | $t_s$ | $\tau_s$ | $\tau_s = t_s\sqrt{\frac{g}{R_{nat}}}$ |
| Double stance time | $t_d$ | $\tau_d$ | $\tau_d = t_d\sqrt{\frac{g}{R_{nat}}}$ |
| Spring stiffness | $K_s$ | $\gamma$ | $\gamma = \frac{K_s R_{nat}}{W}$ |
| Step length | L | $\lambda$ | $\lambda = \frac{L}{R_{nat}}$ |
| Radial Coordinate | R | r | $r = \frac{R}{R_{nat}}$ |
| Initial radial coordinate | $R_0$ | $r_0$ | $r_0 = \frac{R_0}{R_{nat}}$ |
| Radial coordinate at equilibrium | $R_{eq}$ | $r_{eq}$ | $r_{eq} = 1 - \frac{1}{\gamma}$ |
| Radial coordinate at transition | $R_*$ | $r_*$ | $r_* = \frac{R_*}{R_{nat}}$ |
| Height of CoM | H | h | $h = \frac{H}{R_{nat}}$ |
| Height of CoM at mid-stance | $H_0$ | $h_0$ | $h_0 = \frac{H_0}{R_{nat}}$ |
| Spring contraction | $\Delta$ | $\delta$ | $\delta = 1 - r$ |
| Initial spring contraction | $\Delta_0$ | $\delta_0$ | $\delta_0 = 1 - r_0$ |
| Spring contraction at equilibrium | $\Delta_{eq}$ | $\delta_{eq}$ | $\delta_{eq} = \frac{1}{\gamma}$ |
| Spring contraction at transition | $\Delta_*$ | $\delta_*$ | $\delta_* = 1 - r_*$ |
| Angular coordinate | $\theta$ | $\theta$ | N/A |
| Angular coordinate at transition | $\theta_*$ | $\theta_*$ | N/A |
| Initial angular velocity | $\dot{\theta}_0$ | $\Omega_0$ | $\Omega_0 = \dot{\theta}_0\sqrt{\frac{g}{R_{nat}}}$ |
| Maximal Spring Energy | E | $\epsilon$ | $\epsilon = \frac{E}{MgR_{nat}} = \frac{1}{2}\gamma\delta_{\max}^2$ |
| Oscillation frequency | N/A | $\omega$ | $\omega = \sqrt{\gamma}$ |
| Oscillation phase | N/A | $\phi$ | $\phi = \omega t$ |
| Oscillation phase at transition | N/A | $\phi_*$ | N/A |
| Horizontal displacement of the CoM | $X_{com}$ | x | $x = \frac{X_{com}}{R_{nat}}$ |
| Vertical displacement of the CoM | $Y_{com}$ | y | $y = \frac{Y_{com}}{R_{nat}}$ |
| Average CoM's speed | $V_{com}$ | v | $v = \frac{V_{com}}{\sqrt{gR_{nat}}}$ |
| Froude number | N/A | Fr | $Fr = \left(\frac{\lambda}{\tau_s + \tau_d}\right)^2$ |
| Horizontal velocity at transition | N/A | $v_{x*}$ | $v_{x*} \approx (1 - \delta_*)\Omega_0 \cos \theta_*$ |
| Vertical velocity at transition | N/A | $v_{y*}$ | $v_{y*} \approx -(1 - \delta_*)\Omega_0 \sin \theta_*$ |
| Vertical acceleration at transition | N/A | $a_{y*}$ | $a_{y*} \approx \gamma\delta_* \cos \theta_* - 1$ |

### 2.1 Emergence of different walking gaits and their energetics

#### Different gaits are oscillatory modes of the DSLIP model

DSLIP model can function in multiple modes (Geyer et al., 2006; Gan et al., 2018; Andrada et al., 2020; Ding et al., 2022; Mauersberger et al., 2022); these modes include common modes of animal locomotion. These different modes arise from different positions of the CoM in relation to the equilibrium length of the spring (Figure 1). To describe the different modes it is convenient to use the spring compression, Δ ≡ *R* − *R*_*nat*_. Each mode is an oscillation around the fixed point, *R* = *R*_*eq*_ = *R*_*nat*_ − Δ_*eq*_, of the spring-mass system given by the Δ where the spring force balances gravity, Δ_*eq*_ = *Mg*/*K*_*s*_, where *M* is the mass of the subject. Assuming symmetry, at mid-stance the radial coordinate and the height must be either at a maximum or a minimum. At the take-off point, the leg reaches its maximal length or the natural length, *R*_nat_. Whether the mid-stance height is at a maximum or a minimum is determined by the relationship between the compression at mid-stance, Δ_0_, and Δ_*eq*_: If Δ_0_ >Δ_*eq*_, the weight is larger than the spring force at mid-stance, the net vertical force points downwards, the second derivative of the height at mid-stance, 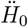,is negative, and the CoM must go down, resulting in a maximum in height and leg length. Thereafter, it must undergo approximately an integral number of oscillations before take-off. Normal human walking with its mid-stance height maxima is the most common gait of this kind with approximately a single radial oscillation between the mid-stance and take-off (Figure 1B, left).

In contrast, if the leg starts below the equilibrium, Δ_0_ < Δ_*eq*_, the spring force is larger than the weight, net vertical force is upward, 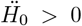,resulting in a height and leg length minima. The radial coordinate undergoes approximately half-integral oscillations before take-off, Figure 1B, middle. The lowest oscillatory mode with approximately half of an oscillation corresponds to the grounded running gait that is employed over a limited speed range in humans but over a large range of speed in some birds ((Andrada et al., 2013b, 2020; Davis et al., 2020). In Figure 1B, right, we also show gait patterns of this type with more than one vertical oscillation.

The gait patterns and the ranges over which they are found, when we have at most one oscillation, are summarized in Figure 2 in non-dimensional units. Due to the centrifugal force resulting from the angular motion, gait transition occurs at a CoM height, *H*_0_, that is slightly higher than the equilibrium height (see Appendix B for a detailed derivation). Due to the centrifugal force, there is a small range of Δ_0_ values for which the gait has a mid-stance maximum in height without an M-shaped GRF (Inverted walking). Finally, there is a large range of values where grounded running, with a height minimum and inverted “U”-shaped vGRF maximum, is observed, consistent with previous work (Andrada et al., 2013b; Blickhan et al., 2018; Andrada et al., 2020). The grounded running and inverted walking gaits are inverted gaits owing to their “U”-shaped vGRF maximum.

**Figure 2.**
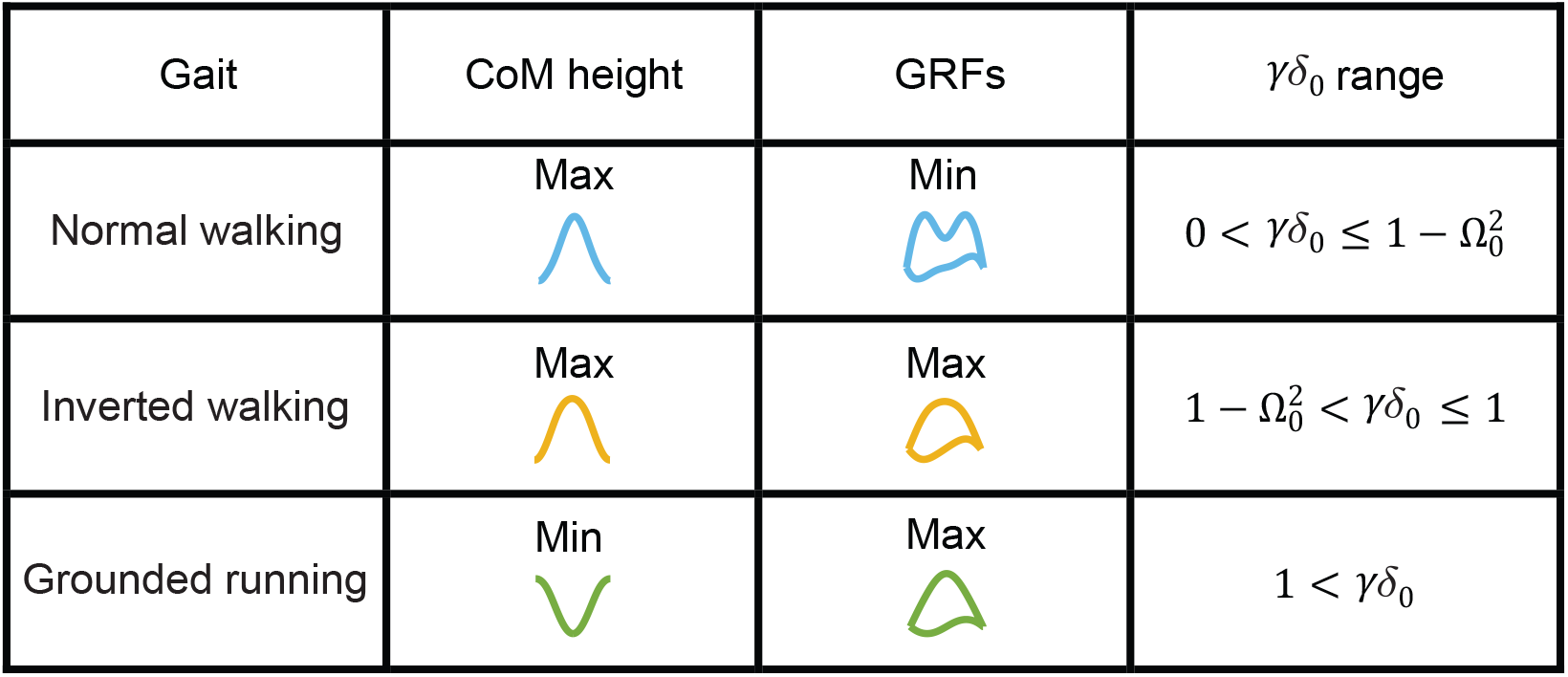
vGRFs and CoM trajectories for different gaits with at most a single contraction-expansion cycle between mid-stance and mid-step and the limits within which each is supposed to occur. The range over which different gaits are observed depends mostly on whether the spring is compressed more or less than the compression necessary to balance the gravitation force. The 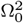 term compensates for the centripetal acceleration and will be small for most walking speeds.

#### Gait parameter space

To evaluate the exact ranges we found limit cycle solutions. A priori, there are five dimensional parameters that control the evolution of a symmetric gait: stiffness and natural length of the leg spring, *K*_*s*_ and *R*_nat_, respectively, the step length, *L*, and the height and angular velocity at midstance, *H*_0_ and 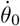,respectively. Time-reversal symmetry requires that at mid-stance and mid-step, 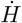 must be zero which provides an additional constraint, leaving only four independent parameters among 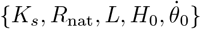 that parametrizes limit cycles. To simplify the analysis further, we used dimensionless quantities (by setting *R*_nat_ = 1): the dimensionless angular speed and length contraction at the mid-stance, Ω_0_ and *δ*_0_, the dimensionless spring constant, *γ*, and relative step length, *λ*. Of these four, only three are independent due to the limit cycle requirement.

The range of speeds, expressed as Froude number, Fr, the square of the dimensionless average velocity (approximately equals Ω_0_^2^), over which limit cycle walking is possible at a given *λ* is shown in Figure 3A. Limit cycles with M-shaped vGRF are found only over part of the speed range over which humans typically walk. Cosistent with previous work, DSLIP cannot model M-shaped walking at the higher end of walking speeds(Geyer, 2005; Geyer et al., 2006; Lipfert et al., 2012; Mauersberger et al., 2022; Lin et al., 2023); this limitation of DSLIP will be explored in the next section. Modes with higher oscillations are found only at low speeds (orange region in Figure 3A). as going through multiple oscillations takes time, increases stance duration, and decreases speed.

**Figure 3.**
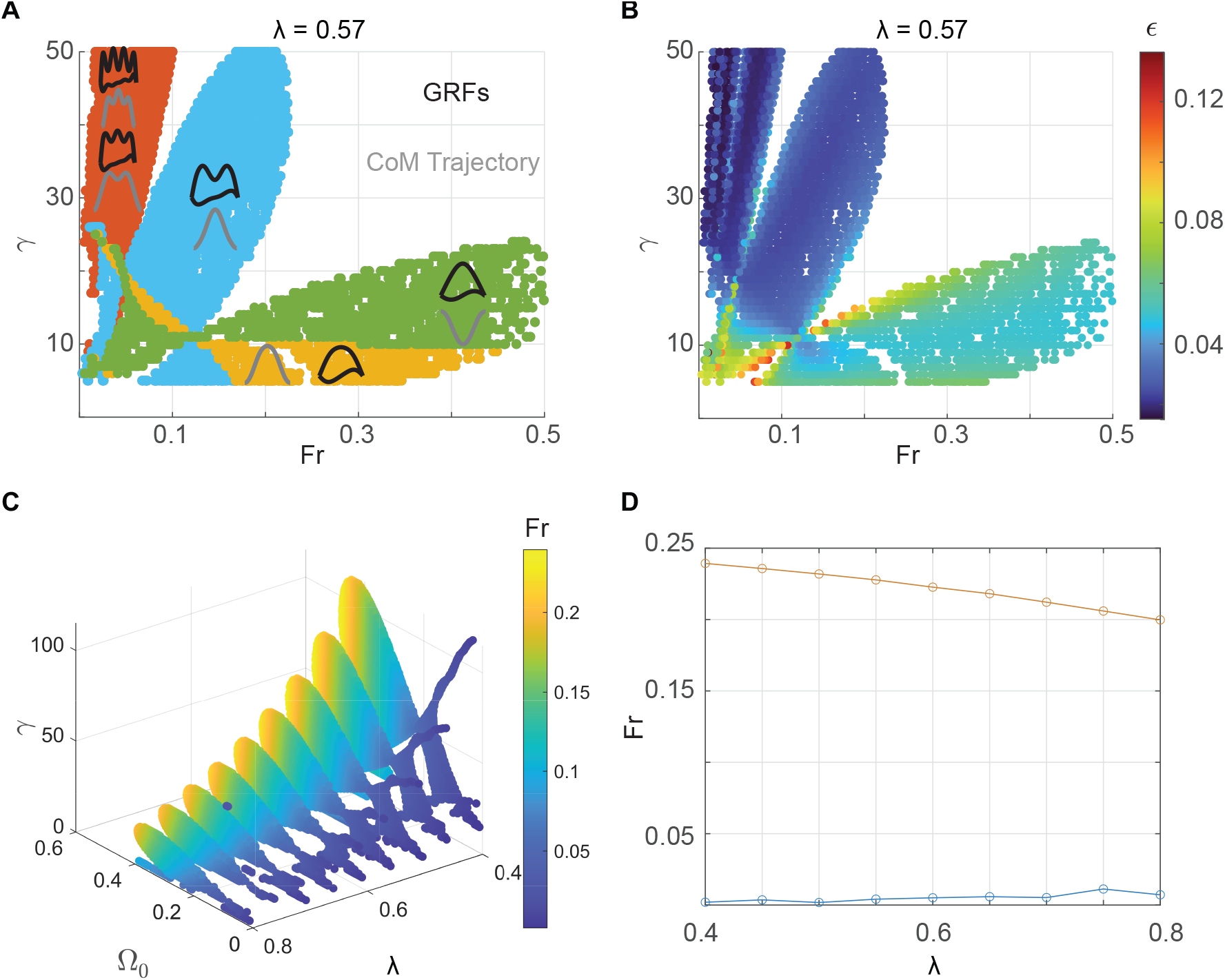
M-shaped walking only occurs only over a limited range of speeds over which it is energetically favored. **A**. Solution space for a fixed value of dimensionless step-length, selected according to the best fit to the experimental data for the preferred walking speed of our subject. Four walking modes are shown - three modes from Figure 2 and one mode with multiple oscillations. The vGRF is shown in black, and the CoM profile is in gray. **B**. The same plot as A, with colors specifying the maximum energy stored in the leg during a cycle shows the M-shaped GRF is the most energy efficient over the range of speeds for walking. **C**. The solution space for M-shaped GRFs for different step lengths. The spring stiffness changes with speed. **D**. The M-shaped walking observed in humans is limited to a Fr of 0.25 across step lengths.

The range of speeds for which a single-humped vGRF (inverted gaits) was observed is more extensive than the M-shaped vGRF. At low speeds, both the M-shaped vGRF and the inverted force profiles are possible using different *γ* values. However, only the inverted force profile is possible at high Froude numbers. Part of this regime (green area) corresponds to grounded running. Consistent with grounded running observed in humans and other bipeds (Andrada et al., 2013b; Blickhan et al., 2018; Andrada et al., 2020; Davis et al., 2020), the spring constant decreases as the gait transitions from normal walking to grounded running.

#### Normal walking gait with “M”-shaped vGRF are preferred because they are energetically efficient

Do humans choose M-shaped GRFs during walking (despite other modes being accessible) because it is energetically favorable? Although DSLIP itself is a conservative model, the spring compression modeled by DSLIP will require work proportional to the energy stored in the SLIP spring. The maximum spring energy stored is a proxy for the energy cost of transport during the given walking step. The maximal stored energy is given by

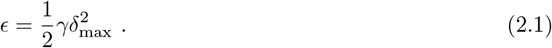

The stored energy for a given walking speed, Ω_0_, for the normal and inverted gaits can be estimated. For the normal gait, *δ*_max_ ≈ 2/*γ* − *δ*_0_, while in the inverted gaits, *δ*_max_ ≈ *δ*_0_ >1/*γ* (Figure 1). In the normal gait, the minimum *ϵ* is achieved by choosing *δ*_0_ → 1/*γ* ⇒ *δ*_max_ → 1/*γ*, so that

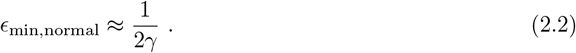

Since *δ*_0_ >1/*γ* in the inverted gaits *ϵ* is minimized as *δ*_0_ → 1/*γ* as well. Note that for the normal gait 1/*γ* is the largest value of *δ*_0_, while for the inverted gait, it is the lowest.

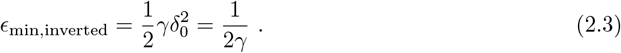

For a given speed, the expression for the minimum stored energy is the same for both gaits, and is inversely proportional to *γ*. Therefore, the gait with higher *γ* - the normal gait Figure 3A - is preferred. The same can be inferred intuitively: The take-off angle, *θ*_off_, does not change much between different walking trajectories. Thus the time, *θ*_off_ /Ω_0_, that a leg is on the ground stays approximately the same as long as the walking speed is the same. However, in this time, during normal walking the radial coordinate must oscillate once for normal walking and only undergoes half an oscillation for grounded walking. Since oscillation frequency goes as the square root of stiffness, *γ*, the normal walking gait must have a larger stiffness.

To quantitatively test this idea, we evaluated *ϵ* throughout the space where we have limit cycle solutions and *ϵ* was smaller for the normal gait compared to the inverted gaits (Figure 3B) for the same speed. Therefore, M-shaped vGRFs are preferable to grounded running because they minimize energy. Gaits with multiple oscillations are even more efficient and should be preferred, but these gaits have a maximum attainable speed; the higher the number of oscillations, the smaller this speed-bound. Within the preferred speed range of human walking, higher oscillatory modes are not available (or have very large stiffness), making the normal walking gait the most energy-efficient gait.

The analysis above ignores the energy used to propel the swing leg; approximate assessment of the energetics of the swing phase show that normal gait will be preferred. It has been previously proposed that the swing energy is ∝ *ν*^4^, where *ν* = 1/(τ_*s*_) is the angular frequency of the swing leg, and τ_*s*_ is the dimensionless time for the single stance/swing phase (Kuo et al., 2005). For a given angular speed, the energy will diminish steeply with *θ*_⋆_ ∝ τ_*s*_. or

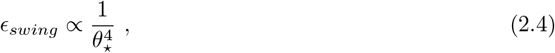

where *θ*_⋆_ is the angular coordinate at the transition from the first single stance to the double stance. For geometrical reasons, just like *θ*_off_, *θ*_⋆_ doesn’t vary much between different gaits, but it does increase slightly (Figure 4B) as one decreases *δ*_⋆_. Since an increase in *γ* decreases *δ*_⋆_, gaits with higher *γ* are preferred.

**Figure 4.**
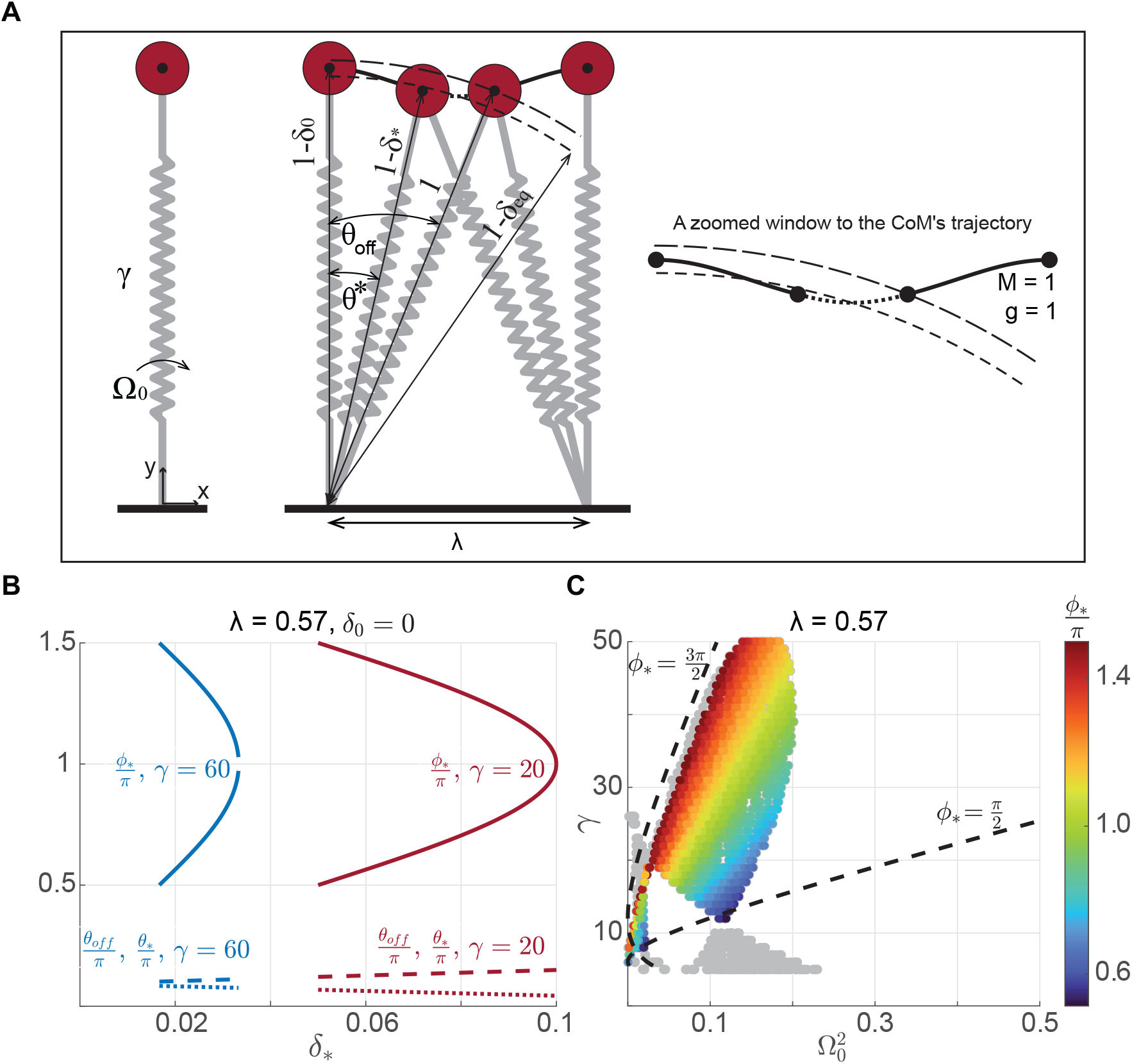
Synchronization between radial and angular motion divides the gaitspace into regions in which different modes are expected. **A**. An example simulation to illustrate synchronization between radial and angular oscillation (middle panel, zoomed version on the right). At a given step-length and leg-contraction at mid-stance, any *γ* and Ω_0_, only solutions that have synchronized radial and angular motion can become a limit cycle. During the time it takes to travel from midstance to the transition between single and double stance phases – denoted by the starred variables, the angular coordinate must go from midstance to *θ*_⋆_. The radial coordinate will go from its position between the natural length and equilibrium length at midstance to a position slightly below it. This corresponds to a change in the value of *ϕ* from 0 at midstance to 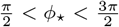 at the transition. **B**. Two examples based on analytical results show that while *ϕ*_⋆_ approximately accesses the entire range defined for normal walking, *θ*_off_ and *θ*_⋆_ slightly increases and decreases respectively. **C**. In the figure, only solutions with 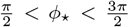 are shown by color bar; the others are gray. Analytical constraints from synchronization are shown by the dashed line, which is close to the lower bound on speed. However, there is no limit on the upper bound. Note that at high *γ*, there are no limit cycle solutions close to 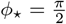.

To investigate the range of speed allowed using the M-shaped GRF pattern, we found the limit cycles for the range of relative step lengths (*λ*) in our experimental data. The region allowed for M-shaped (normal human walking) (Figure 3C) shows that as *λ* decreases, the lowest value of *γ* allowed increases. The maximum and minimum Froude numbers (*Fr*) (Figure 3D) show that DSLIP is a good model at lower speeds but is limited at higher speeds across *λ* values. Compared to the previous study (Antoniak et al., 2019) which assessed the range of Fr numbers allowed using constraints on the single stance, i.e., without any requirement for limit cycles (Antoniak et al., 2019), the allowed speed only is altered at the higher end. Essentially, DSLIP is an adequate model for walking at slow speeds whether one considers just the synchronization of radial and horizontal motions during the single stance or the full gait cycle. In contrast, the range of speed at the high end dramatically decreases when the double stance phase is included, a topic discussed at length in the next section.

### 2.2 Constraints from synchronization of radial and angular motion in single stance and velocity redirection in double stance limit DSLIP normal walking speed

#### Synchronization between radial and angular motion during the single stance describes the lower limits of speed possible with M-shaped GRF

The mechanical constraints that limit the range of speeds for M-shaped walking is not understood. To better understand these constraints, we sought an analytical approximation of the DSLIP model. The analytical approximation has two parts that correspond to single and double stance phases, respectively (see Appendix C for details). First, during the single stance phase, we assume that the angular and radial motion are decoupled. When there is no angular motion, and *θ* ≈ 0, the equation of radial motion can be written as

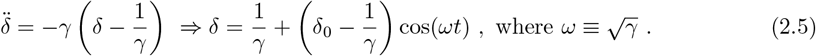

In other words, *δ* simply oscillates around its equilibrium value, 1/*γ*. Further, under the approximation that angular speed is constant, we have

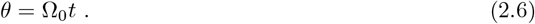

The oscillation phase of the radial motion can be defined as

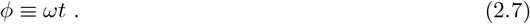

If *ϕ*_⋆_ and *t*_⋆_ denote the oscillatory phase and time when the single stance transitions to the double stance, at this same time the angular motion must traverse up to the transition angle, *θ*_⋆_ (Figure 4A):

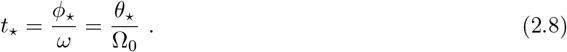

In other words, *γ* and Ω_0_ are related as

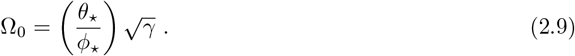

This equation implies that as speed (Ω_0_) increases, the leg must oscillate faster in the radial direction to keep up, leading to a greater stiffness (*γ*). The relationship between (Ω_0_) and (*γ*) is more complex as *θ*_⋆_ and *ϕ*_⋆_ are not constants but rather given by (see Appendix C):

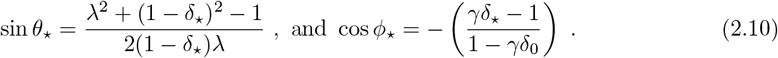

Briefly, the *θ*_⋆_ equation above results from the transition geometry (Figure 4A), and *ϕ*_⋆_ from Eqn. (2.5). Since *δ*_⋆_ is typically small and ranges between 1/*γ* < *δ*_⋆_ < 2/*γ* ≪ 1, *θ*_⋆_ does not change much; there is a small increase with decreasing *δ*_⋆_ (Figure 4B). Assuming *γ* ≫ 1, we have

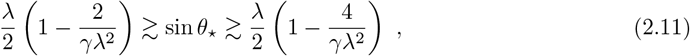

As *γ* increases, *δ*_⋆_ becomes smaller, and accordingly *θ*_⋆_ increases towards sin^−1^(*λ*/2).

In contrast to the small *θ*_⋆_ range, *ϕ*_⋆_ changes considerably (Figure 4B). When cos *ϕ*_⋆_ is negative (we will justify this in the next subsection), *ϕ*_⋆_ can, a priori, take any value in the range

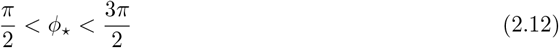

for a single radial oscillation of the CoM. Moreover, (2.10) implies that as *δ*_⋆_ varies, we have two branches of *ϕ*_⋆_(*δ*_⋆_): a branch along which *δ*_⋆_ varies between 1/*γ* to 2/*γ* − *δ*_0_, and *ϕ*_⋆_ varies between *π*/2 to *π*, see Figure 4B and another where *ϕ*_⋆_ goes from *π* to 3*π*/2 as *δ*_⋆_ varies between 2/*γ* − *δ*_0_ to 1/*γ*.

We can estimate the speed bounds based on the analytical equations above. It is clear from (2.9) that Ω_0_ decreases if *ϕ*_⋆_ increases and *θ*_⋆_ decreases, however, the effect of *ϕ*_⋆_ change is much larger. Thus approximately the lower-bound on speed is attained at *δ*_⋆_ → 1/*γ*, and *ϕ*_⋆_ → 3*π*/2 following the upper branch, and yielding

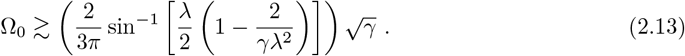

In a similar way, the upper bound is attained as *δ*_⋆_ → 1/*γ*, and *ϕ*_⋆_ → *π*/2:

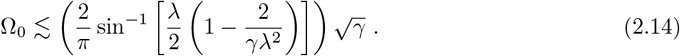

The upper and lower bounds resulting from this synchronization are plotted in Figure 4C. The analytical lower bound derived above matches the simulation results well, implying that the analytical approximation captures the mechanics well. However, the analytical upper bound does not match the bounds obtained through simulation. This mismatch occurs because, except for low *γ*, the allowed *ϕ*_⋆_ does not reach *π*/2; the allowed *ϕ*_⋆_ deviates further from *π*/2 as *γ* increases. Single stance mechanics do not constrain the speed for normal walking; instead, as we will see next, constraints from double support limit *ϕ*_⋆_. This result explains why a previous study that considered only the single-support phase came to the conclusion that DSLIP can function as a model for walking even at high speeds (Antoniak et al., 2019).

#### Limits of DSLIP on speed result from a combination of synchronization and the requirement to redirect vertical velocity component during the double stance phase

The vertical CoM velocity, which is pointed downward at the beginning of the double-support phase, must be redirected upward at the end of the double-support phase (Geyer et al., 2006); the required redirection increases with speed. As speed increases, this redirection becomes more difficult because the double-support phase becomes shorter, and the required change in velocity is larger (Figure 5A): As speed increases, *γ* increases as well, and so does the equilibrium height (1− 1/*γ*). Moreover, as the radial motion of the CoM is approximately oscillating with an amplitude less than 1/*γ*, the CoM trajectory is closer to the natural leg length at higher speeds (Figure 5A), *r* ≲ 1− 2/*γ*. Consequently, the transition geometry dictates that the transition occurs closer to the mid-step at higher speeds. This change, in conjunction with increased horizontal speed, implies that less time is spent in the double support phase. At the same time, as the vertical component of velocity increases with the overall speed increase, a larger change in speed is required at the transition. A larger speed change in a shorter time necessitates a larger acceleration. A back of the envelope calculation is instructive: The double support phase duration, *t*_*d*_ ~ *δθ*/Ω_0_ keeps shrinking as speed increases while the required change of vertical velocity necessary during the double support phase increases, *δv* ~ 2Ω_0_ sin *θ*_⋆_. Thus the average upward acceleration that one needs, 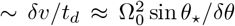,increases with speed. The increased acceleration cannot be produced because the force that can be generated during normal walking is bounded and is equal to the weight (this is the maximum downward force one can have, and in the simple harmonic approximation equals the maximal upward force). Moreover, we will show that in contrast to the synchronization during the single support phase where the highest speeds are attained when *ϕ*_⋆_ is close to *π*/2, a successful velocity redirection is more likely when *ϕ*_⋆_ is close to *π*. This mismatch between the conditions that support the highest velocity in the single and double support phases severely limits the speed bound for M-shaped GRF (Figure 5B). This mismatch is illustrated in (Figure 5B), and only exists for high speeds. The above arguments are qualitative. Below we provide an approximation and provide an analytic condition determining the maximum speed bound for the normal walking gait.

**Figure 5.**
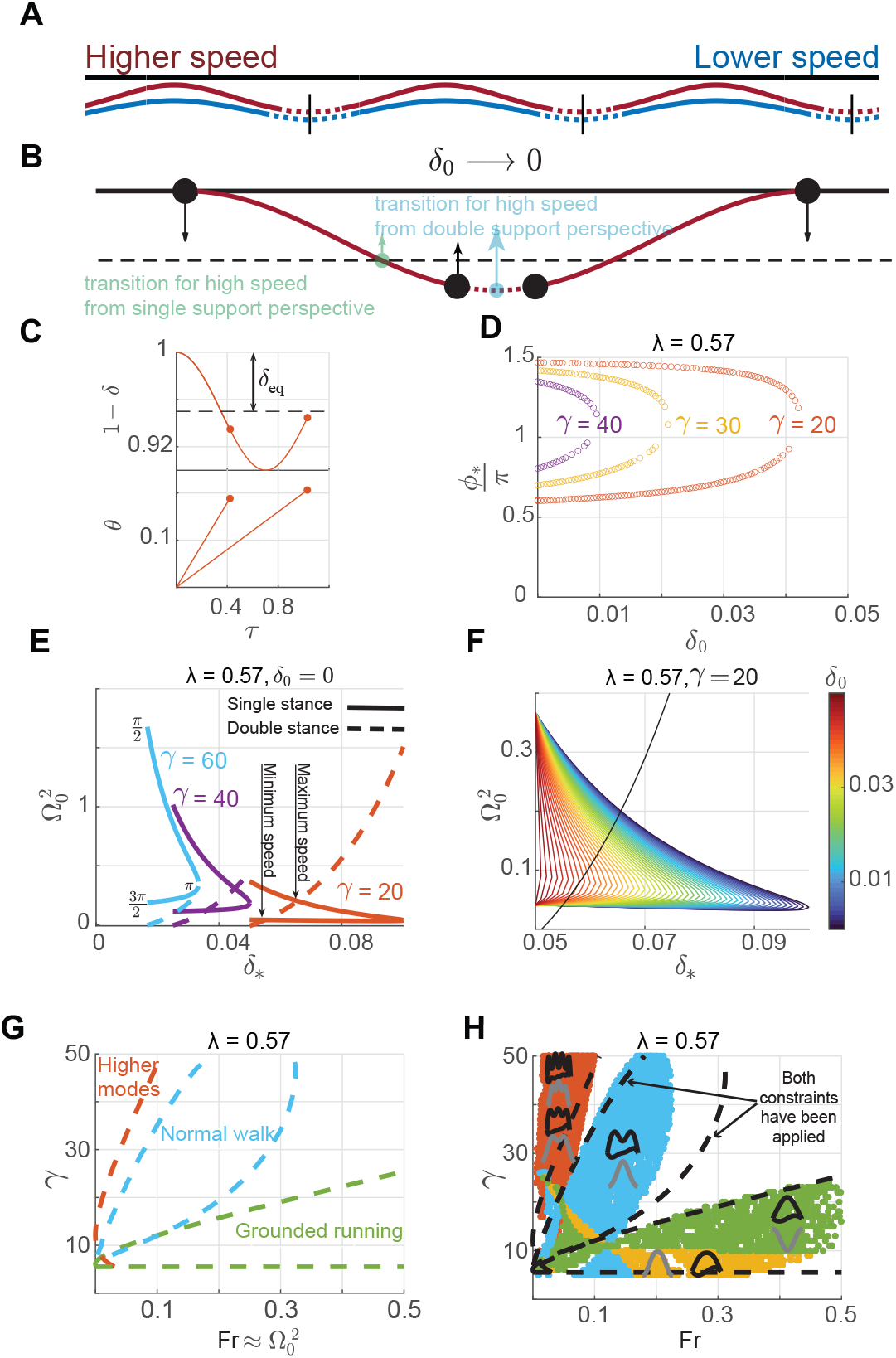
M-shaped walking is limited to low speeds because of a combination of synchronization and velocity redirection constraints. **A**. Two CoM trajectories illustrate the single stance (solid lines) and the double stance (dotted lines) phases. The double stance phase gets shorter with increasing speeds. The vertical black lines specify the mid-step. **B**. The conditions preferred from the single stance synchronization viewpoint (green solid) for high speeds are different from those that will produce the largest redirection forces (cyan). The actual transition point is somewhere between the two extremes. This compromise means that the speeds allowed for M-shaped GRF walking are limited. **C**. The figure shows the approximate evolution of *δ* and *θ* during the single stance phase. There are only two solutions once *γ, δ*_0_, *λ* are fixed. **D**. The variation of *ϕ*_⋆_ vs. *δ*_0_ for a given step-length for the normal walking gait. By increasing *γ*, generally the speed increases, and for a given gamma, as the speed increases *ϕ*_⋆_ and *δ*_0_ values get closer to *π* and zero respectively. **E**. A graphical representation of how single (solid line) and double stance (dashed line) constraints affect the range of possible speeds. Here *δ*_0_ = 0. The highest speed possible (intersection) is much smaller than the highest speed from just single stance considerations(obtained at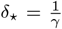). The difference becomes more with higher stiffness until at the highest stiffness (light blue, *γ* = 60), there is no solution (no intersection point). **F**. All solutions for a fixed step-length and stiffness. Note the double stance constraint is independent of *δ*_0_. **G**. The region of different gait patterns that is estimated by our analytical approximation. The boundaries for normal walking become highly constrained. The other boundaries - for grounded running and higher modes - are a result of single stance constraint alone. **H**. Analytical boundaries of walking solutions from G overlayed on the numerical solution for comparison.

To estimate the speed bound we will first derive an approximate analytical solution for limit cycle, and then use this analytical solution to estimate the speed bound. The horizontal speed and vertical acceleration are approximately constant during the double-stance phase and equal to their value at the transition between single and double-stance phases: *v*_*x*_ ≈*v*_*x*⋆_ and *a*_*y*_ ≈*a*_*y*⋆_. In particular, this constancy implies that at the transition, the vertical acceleration must be upward to make velocity redirection possible, or *a*_*y*⋆_ >0. For an approximately simple harmonic radial oscillation, this occurs during the phase, 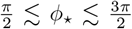, thereby justifying the assumption (2.12) we made earlier. Using these approximations, and the fact that in the time the leg has to travel horizontally to the mid-step from the transition point, the upward force must be sufficient to bring the downward velocity at transition to zero at mid-step, we can derive the relationship Ω_0_(*δ*_0_, *δ*_⋆_, *γ, λ*) (see Appendix D for the derivation) as

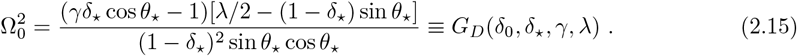

This nonlinear function determining Ω_0_ as a function of *δ*_0_, *δ*_⋆_, *γ, λ* describes the speed based on the double stance constraint. Because *δ*_⋆_ ≪ 1 and *θ*_⋆_ approximately remain a constant, the first term (the net upward force) in the numerator is the most important for determining speed, and this will be important later.

The synchronization relation obtained from the single stance phase is also a function of *δ*_0_, *δ*_⋆_, *γ, λ*:

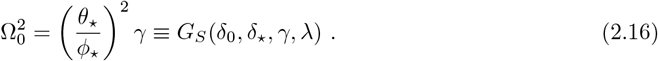

Thus, in order to have a synchronized limit cycle, the four parameters, *δ*_0_, *δ*_⋆_, *γ, λ* must be related,

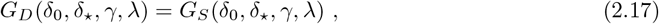

leaving only three independent parameters, *δ*_0_, *γ, λ*. For a given *δ*_0_ and *γ*, inverting the cosine function in (2.10) while obtaining *ϕ*_⋆_ results in two branches, referred to here as *ϕ*_*u*_(*δ*_⋆_) ∈ (*π*, 3*π*/2) and *ϕ*_*l*_(*δ*_⋆_) ∈ (*π*/2, *π*) (Figure 5D). Accordingly, for a given *λ, γ* and *δ*_0_, the upper branch, *ϕ*_*u*_, leads to a branch with lower speeds from the single stance synchronization condition (2.16),

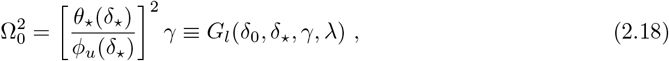

while the lower branch leads to a branch with higher speeds,

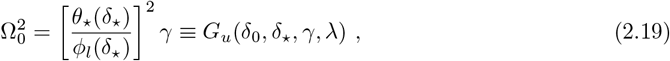

So, if the three parameters, *δ*_0_, *γ, λ* are fixed, there are only two possible values of Ω_0_ resulting from two values of *ϕ*_⋆_ and *δ*_⋆_ corresponding to two branches of solution (Figure 5C); from a different perspective, relating single and double-stance dramatically shrinks the solution space from the entire range between *π*/2 to 3*π*/2 for allowed *ϕ*_⋆_ to just two values of *ϕ*_⋆_ (Figure 5D).

#### Satisfying both single stance and double stance constraints simultaneously is difficult at high speeds and curtails speeds at which walking is possible

Normal walking must satisfy both the single stance and double stance requirement (2.17). The maximum speed occurs at different *δ*_⋆_ and *ϕ*_⋆_ values for the single and double stance: Synchronization during single stance (2.16) suggests that a speed maximum is reached as *δ*_⋆_ → 1/*γ* and *ϕ*_⋆_ → *π*/2 (Figure 5E). However, synchronization during double-stance does not allow *δ*_⋆_ → 1/*γ* and *ϕ*_⋆_ → *π*/2: As *δ*_⋆_ → 1/*γ* - the upward force (the first term within the parenthesis in the numerator of (2.15)) becomes negative and is disallowed (see Figure 4C). Thus, it is not possible for *ϕ*_⋆_ to attain *π*/2 (Figure 5E). This inability of *ϕ*_⋆_ to reach *π*/2 is also reflected in the simulation results in Figure 4C and becomes worse as *γ* increases (Figure 4C and 5D). The maximum upward force in the double stance phase occurs at the largest compression possible, *δ*_⋆_ ≈ 2/*γ* when *ϕ*_⋆_ ≈ *π*. In calculating the force *δ*_⋆_ is multiplied by *γ* ≫ 1, and thus, the effect of *δ*_⋆_ in *G*_*D*_ is dominated by the force term. The maximum speed possible is a compromise between the considerations from single and double stance and the largest speed occurs at a value of *δ*_⋆_ between *π*/2 (where the maximum speed from single stance condition occurs) and *π* (where the maximum speed from double stance condition occurs).

By inspection of (2.10) it is also clear that for a given *δ*_⋆_, *ϕ*_⋆_ is smallest if *δ*_0_ = 0. Thus, the maximum speed is approximately attained at a *δ*_⋆_ that satisfies both (2.16) and (2.15) for *δ*_0_ = 0. Or,

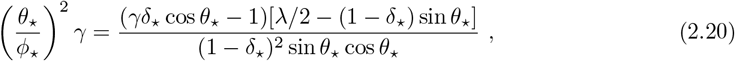

where cos *ϕ*_⋆_ = − (*γδ*_⋆_ −1), and *θ*_⋆_ is given by (2.10). (2.20) can be solved to obtain *δ*_⋆_ as a function of *λ* and *γ*. Graphically, the solution is given as the intersection between curves depicting equations (2.15) and (2.19) or (2.18) (Figure 5E). The constraining function, *G*_*D*_, from the double-support does not depend on *ϕ*_⋆_, and therefore has no branches. It is a monotonically increasing function of *δ*_⋆_ that can intersect both the lower and the higher branches, *G*_*l*_(*δ*_⋆_) and *G*_*u*_(*δ*_⋆_), leading to two possible solutions. The maximum speed is given by the intersection of these two constraints that occur between *ϕ*_⋆_ of *π*/2 and *π*, and is, therefore, lower than the speed possible if we only consider single-stance synchronization. This decrease is exacerbated as *γ* increases (Figure 5E). For a given *λ* and large enough *γ*’s, there are no solutions at all, consistent with our numerical findings (Figure 5E, *γ* = 60). The lower bound is also attained when *δ*_0_ → 0 as that decreases cos *ϕ*_⋆_ so that *ϕ*_⋆_ can get close to 3*π*/2 (Figure 5E). The lower bound is reached when *δ*_⋆_ is close to 1/*γ*, but as argued before, the upper bound *δ*_⋆_ ends up at a compromise value between 1/*γ* and 2/*γ*. The effect of the double-stance constraint on the lower speed bound is much less (Figure 5E).

Essentially the same analysis can be performed for non-zero *δ*_0_ with two limit cycles possible for a given value of *δ*_0_. More generally, the double-valued nature of *ϕ*_⋆_(*δ*_⋆_) leads to a double-valued *δ*_⋆_(*δ*_0_) function (Figure 5F) resulting in a family of curves - one for each *δ*_0_.

The overall results are summarized in Figures 5G and 5H. The single stance constraint alone divides the gait space into contiguous regions with different oscillatory gaits (Figure 4C). Addition of the double stance constraint limits the region allowed (Figure 5G). The results from the analytical approximation of DSLIP and the actual simulations are overlayed in (Figure 5H). The range of speeds predicted from the analytical consideration (see Appendix D for more details) matches the simulation results closely. The correspondence is particularly close for low speeds. The small discrepancy at the higher speed is likely a result of oversimplication of the dynamics of the double stance phase. However, the critical result is that it is the differing constraints from synchronization in the single and double stance phases that limits the range of speed over which M-shaped walking is possible. Two other features of the gaitspace are described in detail in the Appendix F. First, there is a lowerbound on *γ*. Second, the analysis in this section extends to gaits with multiple oscillations.

### 2.3 DSLIP is an adequate model for human walking only for a narrow range of speeds

To evaluate whether the interactions between the walker and the substrate can be quantitatively described with a spring-mass model, we next evaluated how close DSLIP came to describing the kinematics and GRFs during walking. To this end, we fit DSLIP to human walking data. Using an instrumented treadmill, we collected data for four walking speeds - 2.0 mph, 2.5 mph, 3.0 mph, and 3.5 mph (see Supplementary Materials 4.2). Following previous work (Antoniak et al., 2019), we fit the GRF and CoM kinematics in real-world units or dimensional units and individual steps instead of the average data. As choosing the height of the hip marker is a good approximation for the movement of the CoM in time but not the exact CoM location, we began by determining the optimal *R*_*nat*_ for - 2.5 mph - which was the preferred walking speed for the subject (Figure 6A).

**Figure 6.**
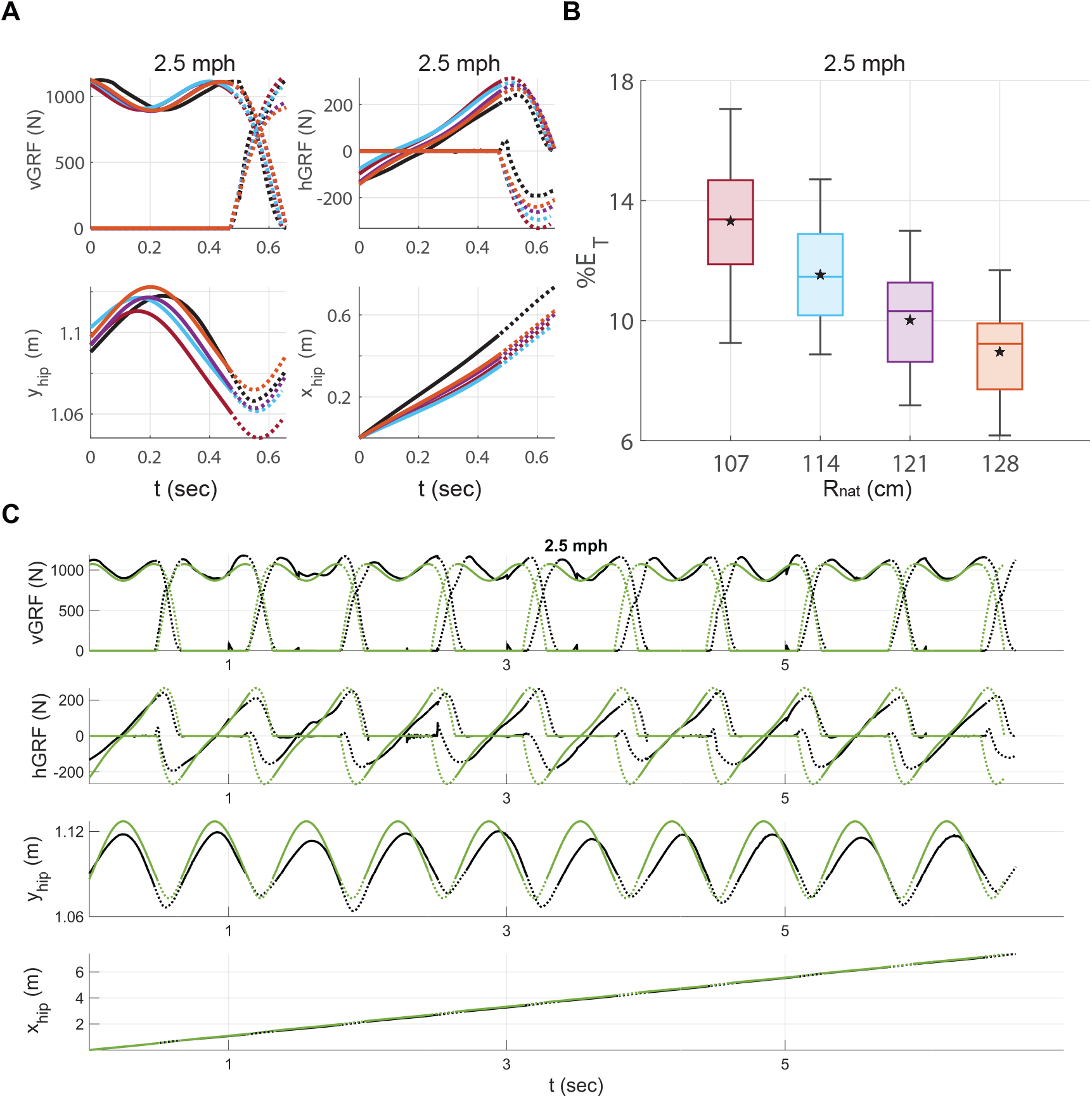
DSLIP is an excellent model for human walking over a narrow range of speeds. **A**. Since the hip marker may not be exactly at the CoM, we fit the experimental data (black lines) to a range of heights both smaller and larger than the hip height (colored lines) (see Supplementary Materials A.1). Solid lines and dotted lines represent single and double stance phases respectively. **B**. The total error for each leg length shows that 128 cm has the lowest error. The error is the sum of errors related to the vGRF, hGRF, height, and horizontal displacement of the CoM. **C**. The optimized limit cycle based on *R*_nat_ = 128*cm* (green lines) fits well into 10 walking steps. The total error in time is negligible.

To this end, we first fit a non-periodic trajectory, i.e., the fits were not constrained to be limit cycles, to each walking step separately, to allow more flexibility and independent assessment of the best fit over 40 steps, thereby increasing statistical power (see Supplementary Materials A.1). In obtaining *R*_*nat*_, we used four values of *R*_*nat*_. The vertical GRF was well fit at all selected values of *R*_*nat*_, as was the height of the CoM. The highest value of *R*_*nat*_, 128 cm, was the best fit to the horizontal GRF (Figure 6A and B), and yielded the lowest overall error, and was selected as *R*_*nat*_ of 128 cm for limit cycle fits.

After fixing *R*_*nat*_, there remained only three free parameters that determine a limit cycle; two of them - the average step length and speed were fixed by constraining them to match the experimentally observed step length and step time. The remaining parameter is selected as the average minimum vGRF over the single stance phase, which can be directly calculated from the data as well.

One example of the limit cycle fit is shown in Figure 6C. A single limit cycle closely describes the entire sequence of steps rather than the average step as is typically done; as a consequence, the limit cycle fits some steps better than others. As an example, the fourth step, which is slower than the optimized limit cycle, does not fit well; but this delay is corrected by faster steps later in the sequence (Figure 6C). Overall, a single limit cycle optimized to fit the entire sequence of steps fits the data well and implies that DSLIP is an excellent model for walking at the preferred speed.

Typical single step fits, one for each of the four speeds, are shown in Figure 7A. Walking at 2.5 mph is best modeled by DSLIP; at this speed, the optimized limit cycle tracks important dynamical features such as the step length, speed, vGRF, and the single stance time (Figure S1D). The model still produces reasonable fits at both 2.0 mph and 3.0 mph, but the fits deteriorate at these speeds. At 2.0 mph, the best-fit model has a longer single stance time; the fitted vGRF oscillates somewhat more than the empirical data. The nature of the deviation is different at 3.0 mph where the model has a lower minimum in vGRF compared to the subject, and much larger vertical CoM oscillations. The model completely fails at 3.5 mph as the minima in the vGRF is close to zero. The average of total errors from GRFs and CoM kinematics along with parameters of optimized limit cycles, are shown in Figure 7B. The total error validates our qualitative observations above showing median errors of < 10 percent at 2.0, 2.5, and 3.0 mph and larger for fits at 3.5 mph.

**Figure 7.**
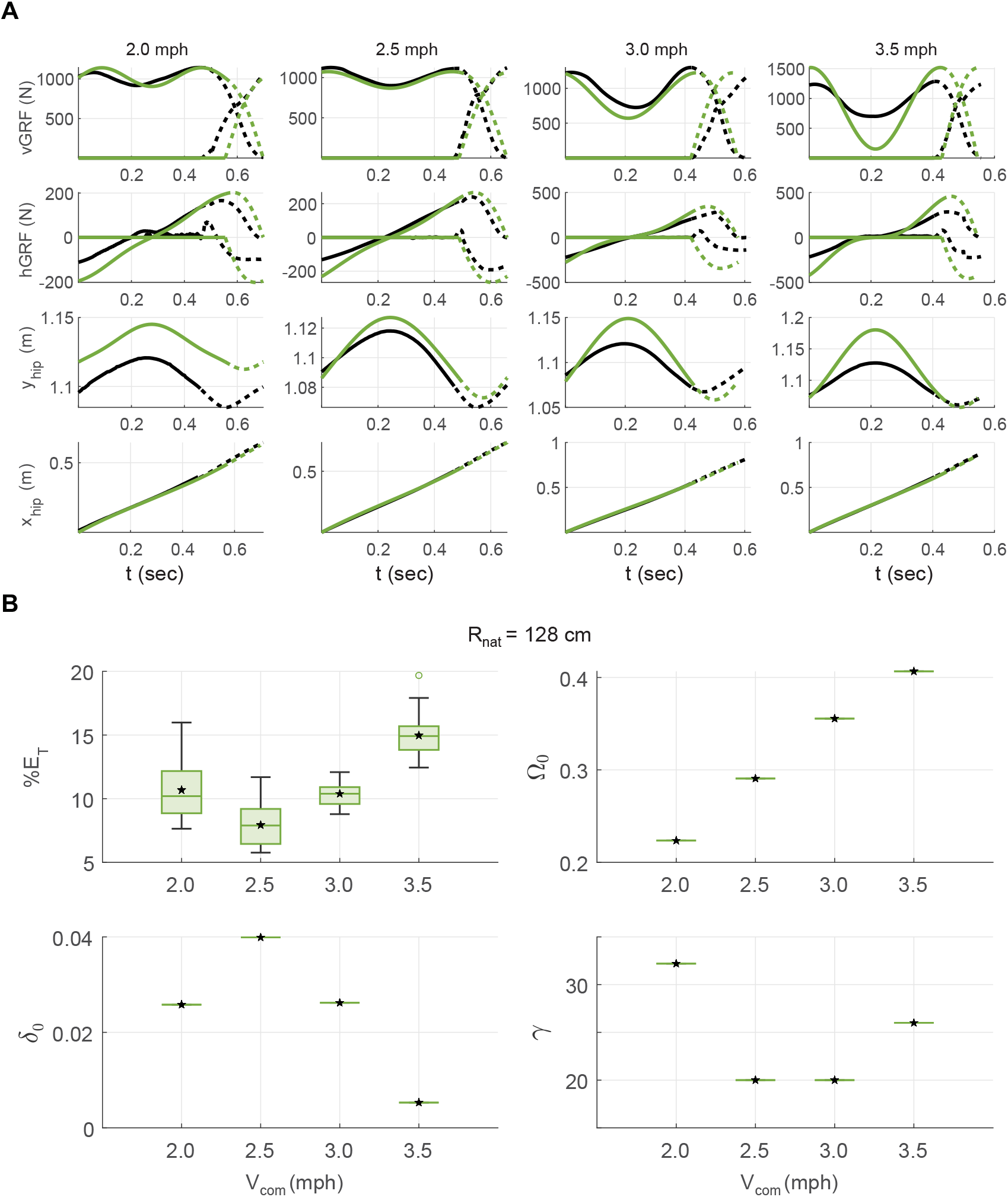
DSLIP fits for both lower and higher than the preferred speed are worse but for distinct reasons. **A**. Example fits (green lines) and data (black lines). Solid lines and dotted lines represent single and double stance phases respectively. The model and subject have the same step length and speed in all fits. We optimized limit cycles based on the values of vGRF at the mid-stance, which can be considered the only free parameter left. The best fit belongs to the preferred speed (2.5 mph), and the highest speed (3.5 mph) has the worst prediction. **B**. The total errors including GRFs and CoM kinematics along with the parameters of the optimized limit cycles for different walking speeds.

Surprisingly, the best-fit spring constant is higher for 2.0 mph (Figure 8A); this finding provides one important clue regarding why DSLIP works as a great model for walking at 2.5 mph and not other speeds. The higher spring constant is unexpected because most previous work has shown that the spring constant decreases as the speed decreases (Farley and Gonzalez, 1996; Kim and Park, 2011). Indeed, the spring constant for the single stance phase, as directly inferred from force-length curve, decreases with speed (Figure S2). At the step length used by our subject to walk at 2.0 mph, there are no limit cycle solutions for this spring constant (Figure 8B) and therefore the spring constant for the best-fit limit cycle is artificially higher. Previous work (Biswas et al., 2018) suggested that at low speeds it becomes increasingly important to model tangential forces. Their introduction may allow one to walk with lower values of *γ* in this low velocity regime and provide a more accurate description of the dynamics. The force-length relationship (Figure S2) also shows that at 2.5 mph, the spring constant during single and double stance phases are similar which explains why a DSLIP model which uses a single spring constant is a quantitative model for human walking at that speed. At higher speeds, the spring constants that describe single-stance and double-stance phases become very different, and this difference makes it difficult for the DSLIP model to describe the data. The fits at 3.0 mph has a stiffness that is intermediate between single and double stance stiffnesses. As a result, the model fit has a smaller *γ* than suggested from the force-length measurements in the single stance phase. With this smaller stiffness, generation of the observed fluctuations in vGRF required a much larger change in the CoM height. In sum, DSLIP seem to function as a quantitative model around the preferred walking speed. At lower speeds, the range of spring constant that can lead to limit cycles shrinks. At higher speeds, the spring constants that describe single and double stance phases are different, making it difficult for DSLIP to model.

**Figure 8.**
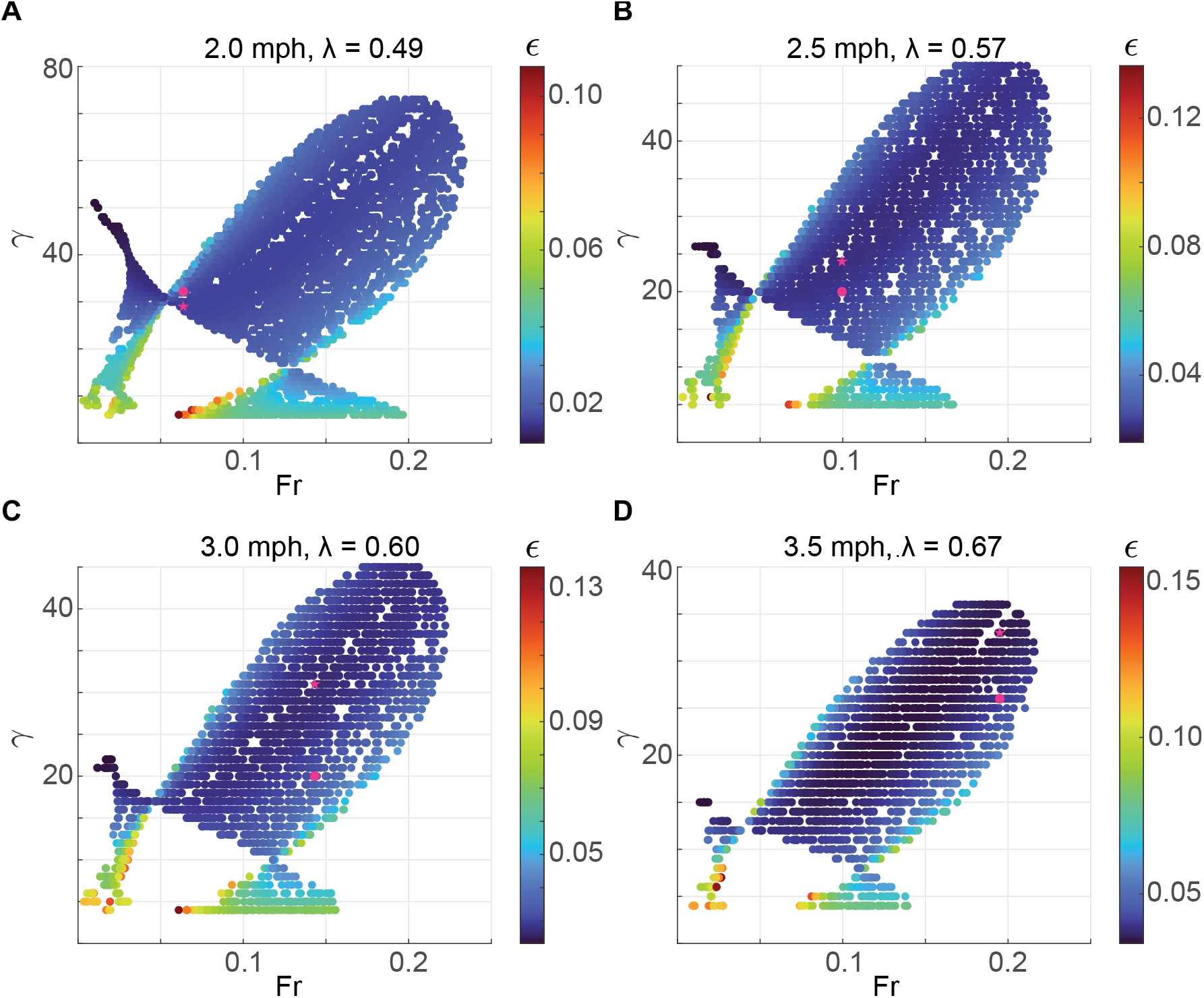
The range of spring constant where a limit cycle is possible likely makes it difficult to obtain good fits for human walking. **A, B, C**, and **D** belong to the 2.0, 2.5, 3.0, and 3.5 mph walking speeds of the subject, respectively. The pink circles show the optimized limit cycles based on our method, and the pink stars show the limit cycles with the minimum energy at the same speed. At both 2 mph and 3.5 mph, the optimization solutions are close to the solution boundary.

## 3 Discussion

### 3.1 A compliant leg is necessary for modeling many features of locomotion

A model with non-compliant legs – IP – continues to persist as a model for walking. The IP model has been successful in explaining the energetics of walking (Donelan et al., 2002; Kuo, 2002; Kuo et al., 2005). The inability of IP model to describe forces is considered a surmountable limitation: by relaxing the impulsive nature of work in the IP model, the model can recover the M-shaped GRF observed during walking. However, we show here that a compliant leg is necessary for modeling many essential features of locomotion including gait choice and speed-dependent gait transitions. First, regarding gait choice, by providing a means to relate leg stiffness that controls the amplitude and period of the vertical oscillation to the angular speed of stance progression, leg compliance provides an analytical framework rooted in mechanics for analyzing which gaits will be observed. In analyzing gait choice, leg compliance is necessary as many of the gaits are not available in the stiff-legged model. We also show here that the energetics of a compliant leg is necessary for understanding why a particular gait, defined by GRF and kinematics, is observed in a given step. Perhaps more importantly, leg compliance is necessary for understanding speed-dependent gait transition as reasons for gait transition in a stiff-limbed walker are completely different from one with compliant legs. It is clear from the analysis performed in this study that the nonholonomic dynamics which results in different optimal transition points for single and double support phases limits the range of speeds over which humans can walk.

### 3.2 M-shaped GRFs are prevalent because they are energetically efficient

An unexplained characteristic of human walking is that humans walk with a M-shaped GRF profile. The M-shaped GRF is observed in other walkers, including both bipeds and quadrupeds (Andrada et al., 2013a, 2014; Basu et al., 2019). At the speeds at which humans walk, other modes of walking, such as grounded running, are possible. However, the M-shaped profile is energetically favored. We have shown that the normal walking gait has a stiffer leg as compared to grounded running, which is preferred because a stiff leg results in smaller vertical oscillations and therefore ultimately less work. This same logic would posit that even higher modes of oscillation with even stiffer legs would be more energy efficient than the normal gait. Accordingly, we observe multi-oscillatory gaits at low speeds (Figure S3). these gaits are not available at typical walking speeds. The unavailability of higher oscillatory modes together with the energy efficiency argument selects the single oscillatory normal walking gait with the characteristic “M”-shaped vGRF for mammals.

### 3.2 Gait transition occurs because velocity redirection is difficult

An important issue that has received much attention is gait transitions: at what speeds do they happen and why? One approach to this problem is using the IP model. Walking using an IP model is not possible at high speeds because at high speeds – above Fr of 1 - the centrifugal force cannot be canceled by gravitational force. This logic was later modified to take into account the fact that the vertical component of the gravitational force would be lowest near the end of the step (Usherwood, 2005) which predicts a transition speed of Fr ~ 0.5. This argument becomes invalid in compliant legs which allows part of the centrifugal forces to provide radial acceleration. Moreover, in transitions with double stance phases, the leg take-off is not a problem as transitions are between double and single stance. Analysis in this study using the DSLIP model comes to a different conclusion for gait transitions. First, even if we take a nuanced approach to walking and impose the condition that walking must have a vGRF minimum at mid-stance, centripetal force does not pose a stringent constraint (see Figure 4C). Moreover, DSLIP makes it possible to walk with gaits that are not possible using the IP model such as the grounded running gait. In sum, adding compliance to the leg removes the appearance of unphysical negative tension force as a reason for gait transition.

The reason for gait transitions in a compliant walker is completely different, and (to us) highly non-intutive, and involves three factors: The existence of a vGRF minimum at mid-stance (a defining feature of normal walking), synchronization of horizontal and vertical motions during the single stance, and velocity redirection in the double stance. The single stance synchronization implies that the highest speed at any given spring stiffness occurs when the single support to double support transition occurs at a different phase (in the oscillatory walking cycle) compared to the phase preferred for the velocity redirection. This tension between single stance synchronization and double stance velocity redirection is further accentuated by the requirement of vGRF minimum at midstance which effectively limits the maximal vertical force the legs can exert during double support to affect velocity redirection. It is this tension between the constraints from the single and double support phases that result in M-shaped GRFs being impossible as a gait at high walking speeds. There are two options when transitioning from M-shaped walking: Transition can be to other walking modes such as grounded running and inverted walking or to running with an aerial phase. Thus, analysis using DSLIP model suggests two different answers to gait transitions: Transitions out of M-shaped GRFs occur at low speeds, but transitions from locomotion without an aerial phase to one with an aerial phase can occur at any speed. Both grounded running and aerial running can occur over a large range of speeds.

At what speed aerial running is preferred depends on the individual and species. In humans, transitions can occur from M-shaped walking to aerial running as is suggested by some. Under certain conditions, there is a small range of speed over which humans walk with a grounded running gait (Shorten and Pisciotta, 2017; Bonnaerens et al., 2019). In many birds, grounded running is preferred over a large range of speeds, often exceeding a Fr of 1 (Andrada et al., 2020). Many non-human primates also prefer grounded running (Blickhan et al., 2018). Fast-running insects and spiders prefer grounded running (Reinhardt and Blickhan, 2014). To address which gait is preferred energy estimates for aerial and grounded running at a given speed must be made, which is beyond the scope of this paper.

### 3.4 Limitations of DSLIP and how they might be overcome

DSLIP is a great conceptual model, but with its simplicity comes some limitations. Among them is the fact that DSLIP cannot support M-shaped vGRF walking beyond 0.25 whereas humans can walk with M-shaped vGRF upto a Fr number of 0.45 (Kram et al., 1997). There are many mechanisms that might contribute to humans walking at higher Fr numbers. One mechanism is that human legs are not massless, and recoil from the leg swinging forward contributes to velocity redirection (Adamczyk and Kuo, 2009). Another mechanism is that the center of pressure moves forward during stance; this forward movement might increase the range of speeds.

All of these processes can be modeled as additions to the DSLIP model and aspects of these processes have been explored by others (Whittington and Thelen, 2009; Lim and Park, 2019; Mauersberger et al., 2022). Adding features to the model will increase model complexity; the two additions below can be highly beneficial without increasing model complexity. One addition is to use a variable spring stiffness. Both the spring constant and the natural leg length during the single and double stance phases are different at high walking speed (Figure S2). This difference suggests that changing the stiffness and natural length of the spring during the double stance phase may be a mechanism for increasing the speed over which M-shaped GRF walking gaits are possible.

Another mechanism is adding an angular spring. As has been noted previously, net forces during walking do not point along the leg but are more vertical (Maus et al., 2010; Müller et al., 2017; Antoniak et al., 2019). This limitation can be addressed by adding an angular spring as we have proposed earlier (Biswas et al., 2018; Antoniak et al., 2019). An angular spring produces restorative forces such that there is no angular force at mid-stance. The angular forces increase as the leg moves away from mid-stance. As investigated in (Biswas et al., 2018), such angular forces can provide a much wider range of realistic gaits at low speeds.

## 4 Material and Methods

In this section, we briefly describe the model, the essential details related to the empirical data, and the numerical techniques to find walking solutions and optimized trajectories. Details are in Supplementary materials.

### 4.1 Walking dynamics of DSLIP

The model is the same as the one introduced by Geyer et al. (Geyer et al., 2006). It has two degrees of freedom (DoF) that describes the sagittal plan motion of a point mass under gravity and spring forces.

#### 4.1.1 The equations of motion

The model in its full dimension and dimensionless form is shown in Figure 1A and Figure 4A respectively. Figure 1A is a schematic but Figure 4A is based on simulation. The model consists of two massless springy legs hinged with a large mass, M, at the hip (CoM). The model does not include any swing phase dynamics, so the single stance phase is described by just a single spring with the mass at the top. The natural leg length of the springs is denoted by *R*_*nat*_. The leg stiffness, *K*_*s*_, and the step length, *L*, are made dimensionless according to the following equations:

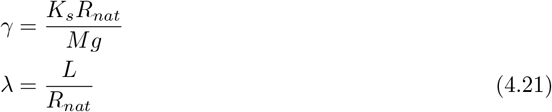

where *g* is the gravitational acceleration. The dynamics during the single stance phase evolve according to the following equations represented in the Cartesian form:

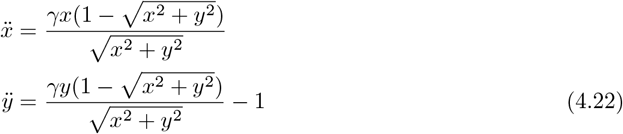

where *x* and *y* denote the dimensionless form of horizontal and vertical displacement of the CoM respectively:

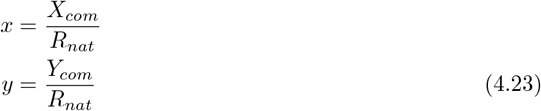

Also, we made the time dimensionless by defining

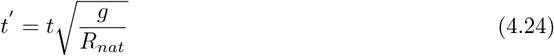

Then, the initial conditions are specified by the position and velocity of the CoM at the mid-stance:

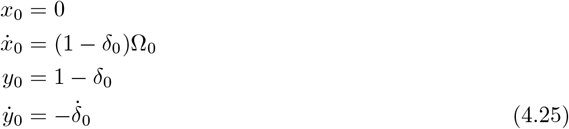

where *δ*_0_ and Ω_0_ are the initial dimensionless spring contraction and angular velocity at the midstance, respectively. Touchdown occurs at a predefined step length. At this moment, the following algebraic equation is satisfied by the CoM position:

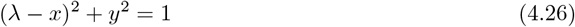

Following touchdown, both the velocity and acceleration of the swing foot becomes zero, and the governing equations changes as follows:

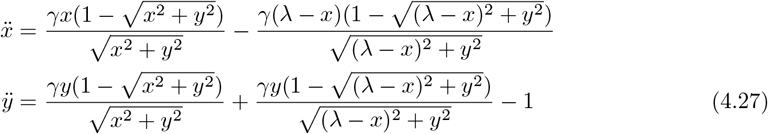

When the contact force at the trailing leg becomes zero, the leg reaches its natural length and leaves the ground. This moment is called toe-off and is defined by:

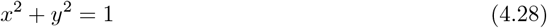

Then the single stance phase restarts by resetting the origin of the coordinate system to the new contact point. In this regard, despite the CoM’s motion being continuous, its *x*-coordinate experiences a discontinuity due to the origin shift:

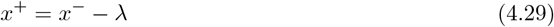

where *x*^+^ and *x*^−^ are the x-coordinates of CoM just after toe-off and before it respectively. The gait cycle ends when the stance leg re-stands vertically (*x* = 0). Now, we can summarize all equations in a single Poincaré return map which maps the states from *i*^*th*^ mid-stance to (*i* + 1)^*th*^ mid-stance:

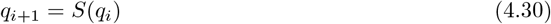

where:

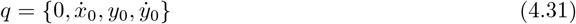

At a fixed point which represents a limit cycle we have:

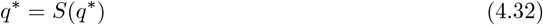

where:

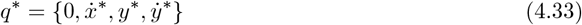

#### 4.1.2 Parameters and Conditions for symmetric human-like limit cycle walking

We focus on symmetric limit cycle solutions. To this end, the first derivative of vGRF must be zero at mid-stance. So we have:

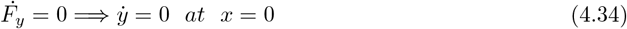

As a result, the general form of initial conditions for equations (4.22) will be:

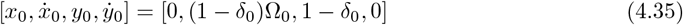

For limit cycles, there is a relation between *δ*_0_ and Ω_0_ to synchronize the radial displacement of the spring with its rotational movement reducing the number of free parameters to be any three of the following four parameters:

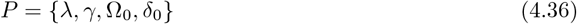

In limit cycles with M-shaped vGRF and a maximum in height at the mid-stance, another constraint was applied resulting in the following constraint 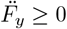 and 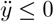 leading to the following inequality:

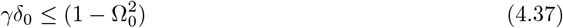

Using these equations and constraints, we found the limit cycles using standard techniques (see Appendix A.1).

### 4.2 Collection of walking data and fitting DSLIP to walking data

The experimental data is collected from walking of a healthy subject (111 kg weight, 185 cm height) on a treadmill for one hundred steps at five different speeds, ranging from 1.5 to 3.5 mph, in increments of 0.5 mph representing the slow, normal, and fast walking of the subject. It is obtained based on the self-selected speed of the subject, followed by 20% and 40% slower and faster speeds. The GRFs were measured by force plates at 1000 Hz, and the hip coordinates were sampled by VICON at 200 Hz. Due to a high level of noise, we excluded data related to 1.5 mph from our analysis. The data was smoothed using the ’smoothdata’ function in MATLAB, using the ’sgolay’ method (Savitzky-Golay filter) which employs a quadratic polynomial fit to smooth data.

To assess DSLIP as a model for human walking, we employed two different strategies. First, we fit the model to each step separately giving us an individual non-periodic trajectory for each step (optimized non-periodic trajectory). Second, by averaging empirical data for each walking speed, we fit a single limit cycle to all steps. The methods used to fit the walking data are described in supplementary methods (see Appendix A.2).

## Acknowledgements

T.B. was funded by the Howard Hughes Medical Institute. This research was supported by RO1DC015827 (VB), RO1NS097881 (VB) and an NSF CAREER award (IOS-1652647 to VB).

## Supplementary Materials

### A Fitting limit cycles to human data

#### A.1 Finding Limit cycles

In general, finding limit cycles is not easy; especially for unstable trajectories. If the DoF is low, and if we have a good estimation of initial conditions, it is easier to find them. Based on trial and error or using analytical approximations, we can find such an estimation around the desired fixed point which represents a limit cycle. At a fixed point we have:

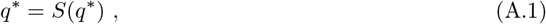

where

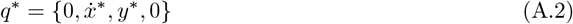

This fixed point represents the initial condition that leads to a symmetric periodic gait. To find the fixed point and analyze its stability, we engage a method described in (Wisse et al., 2004). To explain the method in detail, a small perturbation is added to the fixed point at step *i* as follows:

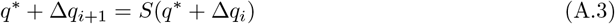

Now, by using the Taylor expansion of Poincaré map around the fixed point we have,

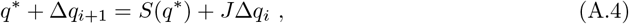

which results in,

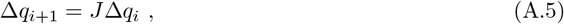

where *J* is defined as,

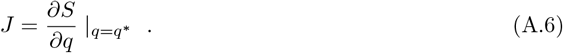

On the other hand, we have,

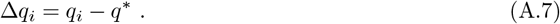

Afterward, based on Eqs. (A.5) and (A.7), we can conclude,

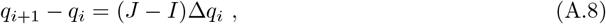

where *I* is the identity matrix. Also, employing Eq. (4.30) leads to

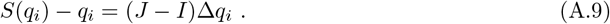

Next, a computer program could be written based on the following algorithm:

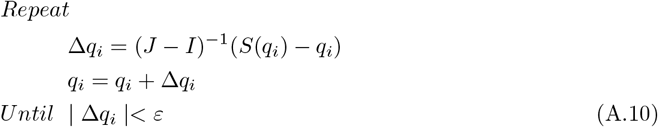

where ε is a small disturbance added to the system. Also, the Jacobian matrix, *J*, can be numerically calculated in every iteration. Now, if the algorithm is convergent, the fixed point and its corresponding Jacobian matrix simultaneously emerge. Otherwise, either the algorithm must be run again with a new initial guess or we need to change the system’s parameters. Finding the first fixed point would help to find other fixed points in its vicinity. In this regard, a new initial guess is defined as a point near the found fixed point. Therefore, trial-and-error is merely necessary to reveal the first limit cycle.

#### A.2 Fitting DSLIP to human walking data

This section describes methods used to fit human walking data without requiring these fits to be limit cycles. With symmetry and limit cycles constraints relaxed, the increased parameter set could lead to a better fit. There are two parameters pertaining to the model, *R*_*nat*_, and *K*_*s*_, the natural length and stiffness of the effective leg, and four parameters pertaining to the initial conditions, the coordinates and their derivatives, 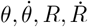.Now both for computational purposes and for linking the data to our theoretical gait analysis, it is convenient to use dimensionless parameters while performing the numerical fits. A key parameter that connects dimensional and dimensionless quantities is *R*_*nat*_, and unlike the other parameters, for biological reasons discussed in the main paper, we expect this to be close to the observed leg lengths and remain the same over all speeds and steps. Thus, our first goal was to determine an appropriate value of *R*_*nat*_ that we will fix for all subsequent fits.

There are two unknowns that we need to consider while estimating *R*_*nat*_. First, because the exact location of the CoM is unknown but related to the hip position, a new parameter, −10 *cm*≤ *D* ≤ 10 *cm*, which defines the vertical distance between the hip and CoM was introduced; when *D* >0, the CoM is over the hip. Since the average mid-stance hip height was measured to be ≈ 112 *cm*, this constrained the CoM height at mid stance, *R*_0_, to lie within, 102 *cm* ≤ *R*_0_ ≤ 122 *cm*. Estimating the amount of spring contraction at mid-stance to be around 5% (or *R*_*nat*_ = 1.05*R*_0_), we therefore constrained *R*_*nat*_ to lie within the range,

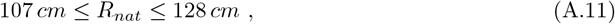

rather than allowing it to take arbitrary values which, as we have discussed in the main paper, can lead to biologically inconsistent results/fits. Based on the range, we next chose 4 different values for *R*_*nat*_: 107 cm, 114 cm, 121 cm, 128 cm, and performed model fits to the experimental data. In a preliminary analysis, we found that minimizing over the entire step (*i*.*e*. over all time points) made it difficult to find the global minimum. To avoid local minimums as well as keep the important features of human walking, a cost function to minimize the errors in the following quantities were chose, minimum of vGRF, peaks of vGRF, CoM height corresponding to the minimum of vGRF, single stance time, step-time, and step-length. Specifically, we considered the cost function *C* to be a sum of differences between the experimental and model trajectories of all the listed quantities in their dimensionless forms:

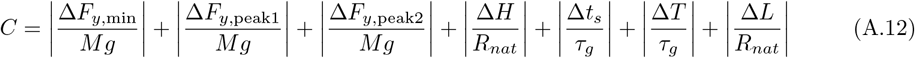

and minimized it over the five remaining parameters. Here we have defined 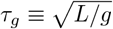. The ’Global Optimization Toolbox’ of MATLAB along with ’fmincon’ function, ’sqp’ algorithm, and ’MultiStart’ object was employed. The bounds on the stiffness and initial conditions were:

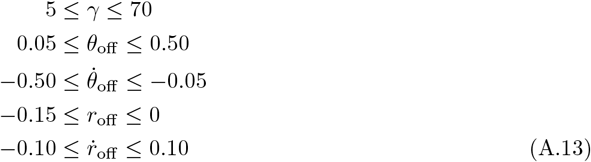

where index ‘off’ indicates the moment of toe-off. We selected the toe-off moment rather than mid-stance because detecting mid-stance is considerably more challenging due to the inherently asymmetrical nature of walking. In contrast, the toe-off moment can be easily identified from data irrespective of symmetry conditions. Based on this rationale, we found optimized fits for the specified values of *R*_*nat*_ for all the steps with four different walking speeds, ranging from 2.0 mph to 3.5 mph, in increments of 0.5 mph. Then we computed the root-mean-square error (*RMSE*), for normalized hGRF, vGRF, CoM horizontal displacement, and CoM height:

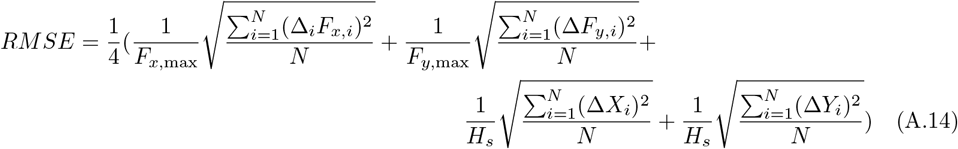

where *H*_*s*_ is the height of the subject. *RMSE* was calculated for all the time points, *N*, and we found the total error coming from all the steps for each value of *R*_*nat*_; *R*_*nat*_ = 128 cm gave the smallest error. Therefore this value was chosen to find the optimized limit cycle fits, as we now discuss.

#### A.3 Optimized limit cycles

Since limit cycles describe a harmonic motion in a dynamical system, it is not meaningful to look for them for each step separately. In contrast, by considering human walking as a dynamical system, it is more acceptable to fit a single limit cycle to all steps. To this end, first, we fixed the values of *R*_*nat*_ and *D* to the values obtained in the previous section from fitting the human trajectories. Next, *λ* was chosen based on the average step length of the subject for each speed divided by *R*_*nat*_. The next parameter was selected to be the subject’s average speed or equivalently the Froude number, *Fr*, which was calculated according to,

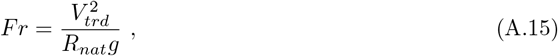

where, *V*_*trd*_, is the treadmill speed. Thus only one parameter was left to be determined to have a full dimension limit cycle, emulating the GRFs and CoM’s trajectory of the subject. Here, we chose the dimensionless form of vGRF at the mid-stance (which approximately corresponds to the minimum vGRF of the subject during the single stance phase divided by the weight of the subject), *γδ*_0_, since it was available from the data. The other choice could have beeen the single stance time; however because we already had the step length and the speed of the subject, the period of the cycle was fixed. So instead of tracking another kinematic variable, we reasoned that it would be better to try to fit a quantity that captured the force profile. By plotting the solution space of limit cycles for the fixed *λ* in the *Fr*-*γδ*_0_ plane, we were able to determine the limit cycle which has the same *λ* and *Fr* as the subject, and had a value of minimum vGRF (*γδ*_0_) that was closest to the empirical data. The difference between the model and the emperical value also provided an effective way to judge the model accuracy.

### B Single stance dynamics and different gait patterns

Let us characterize the different gaits DSLIP can realize that, at most, exhibit a single radial (leg length) oscillation. We can write the equations of motion for the single stance phase in dimensionless polar coordinates (Table 1) centered around the point of ground contact as

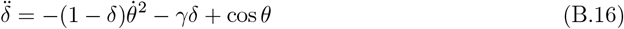

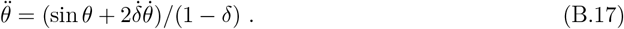

where *δ* represents the dimensionless spring contraction, and *θ* defines the angular coordinate.

For the most part we are going to consider symmetric limit cycles. This means that at the midstance the CoM either has a minimum or a maximum in kinematic variables such as vertical height, and vGRF. We will show that three different gaits are possible depending upon the height and vGRF profiles with at most a single radial (leg length) oscillation. The normal gait has a height maximum and vGRF minimum at mid-stance, while both the inverted gaits (grounded running and inverted walking) have a vGRF maximum at mid-stance. While the grounded running has a height minimum at mid-stance, inverted walking has a height maximum similar to normal walking gait. Finally, let us reiterate (Biswas et al., 2018) that within the DSLIP model there is no provision to have a midstance maximum in horizontal velocity, it always has a minimum. The different gait characteristics are summarized in the table below along with the relationships between gait parameters that must be satisfied. We now derive these relationships.

To ascertain the region in parameter space where the different gaits emerge, first consider the vertical acceleration at the mid-stance:

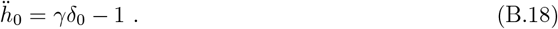

Clearly then to have a maximum in height we must have *δ*_0_ < 1/*γ*. Now the vertical spring force is given by

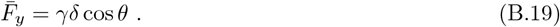

Using (B.17) we find that at the mid-stance

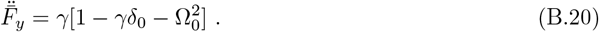

For 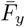 to have a minimum at the mid-stance then, this must be positive, or

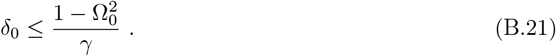

We also note that for a given *λ* there is an upperbound for *δ*_0_ to have any single stance phase at all, see Fig.?:

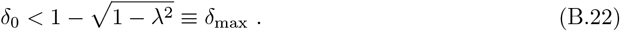

Thus based on the range of *δ*_0_ one can have different gait profiles that we tabulate below:

| gait | mid-stance height | mid-step height | mid-stance vGRF | mid-step vGRF | mid-stance velocity | $\delta_0$ range |
| --- | --- | --- | --- | --- | --- | --- |
| normal walk | maximum | minimum | minimum | maximum | minimum | $0 < \delta_0 < \frac{1 - \Omega_0^2}{\gamma}$ |
| inverted walk | maximum | minimum | maximum | minimum | minimum | $\frac{1 - \Omega_0^2}{\gamma} < \delta_0 < \frac{1}{\gamma}$ |
| grounded run | minimum | maximum | maximum | minimum | minimum | $\frac{1}{\gamma} < \delta_0$ |

### C Approximate Trajectories

Our goal in this subsection is to derive approximate trajectories of the CoM in order to gain analytical insights into how different parameters must adjust to have a synchronized motion. Also, this will help us address how well DSLIP is able to capture some of the well-known features of the walking gait.

#### C.1 Single stance phase

Starting from the dynamical equations in polar coordinates (B.16, B.17), solutions for *δ*(*t*) and *ϕ*(*t*) were derived in the main manuscript,

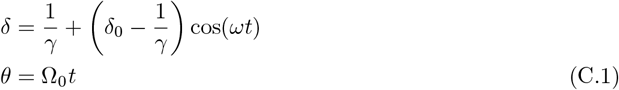

where we assumed that the angular and radial motion during the single stance phase are effectively decoupled. The main idea behind this approximation is that for walking trajectories the radial motion undergoes oscillations around its equilibrium position *δ*_*eq*_ = 1/*γ*, and since *γ* ~ 𝒪 (10) − 𝒪 (100), the oscillations are small. We also assume that the angular/horizontal velocity of CoM remains approximately constant. Technically, this means that we are ignoring 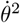 term as compared to (*γδ*) in (B.16). Since 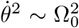 and (*γδ*) ~ 1 on an average, this boils down to assuming 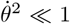 which is valid for the speeds we are interested in. We also assume that 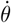 is approximately constant, or 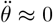.By inspection of (B.17) 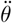 depends on *θ* but this is small, *θ* < *λ*/2, for the steplengths under consideration. 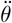 also depends on 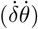.While 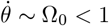,on an average 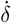 is close to zero suggesting a small effect coming from this term 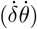.We shall see, that these approximations provide valuable qualitative and quantitative insights into the dynamics and the relationship between various relevant dynamical parameters.

#### C.2 Transition to double stance

As argued in the main manuscript, synchronization between the radial and angular motion relates *γ* and Ω_0_ as

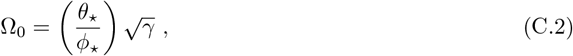

where ⋆ marks the values at the transition point between the single and double stance phases. From geometry, we can find *θ*_⋆_ as

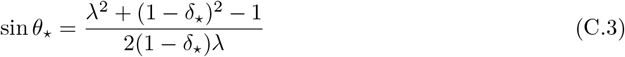

while substituting *δ* = *δ*_⋆_, and *ωt* = *ϕ*_⋆_ in (C.1) yields

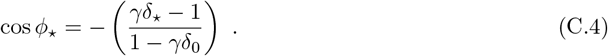

The dependence of *θ*_⋆_, *ϕ*_⋆_ as a function of *δ*_⋆_ are plotted in Figure 4B. In principle, the transition time can be found by solving

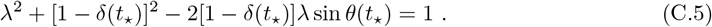

so that *t*_⋆_ = *t*_⋆_(*λ*, Ω_0_, *γ, δ*_0_). One can then evaluate *θ*_⋆_ = *θ*(*t*_⋆_), and *δ*_⋆_ = *δ*(*t*_⋆_), to obtain the position of CoM at the transition, as well as the phase angle, *ϕ*_⋆_ = *ωt*_⋆_. Thus all these quantities can be thought of as functions of four gait parameters, *λ*, Ω_0_, *γ*, and *δ*_0_.

#### C.3 Double stance phase

To approximate the double stance phase we are going to assume that the horizontal velocity and the vertical acceleration remain approximately constant. The intuition behind these approximations is as follows: the two springy legs provide horizontal forces in opposing directions so that we expect the average horizontal acceleration to be small and therefore the horizontal velocity to remain approximately constant. Realistic walking trajectories typically exhibit low-velocity changes which further strengthens this argument, and we compute the horizontal velocity at the start of the double stance. In contrast with the horizontal motion where the legs oppose each other, both legs provide a vertically upward forces. At the beginning of the double stance phase, all the force comes from the leg that was supporting the single stance phase at the touchdown, the swing leg is at its natural length. Thereafter, while the initial stance leg unloads, the leg that touched down loads. Therefore we conjectured that the net upward force may not change much, and approximate the net force as a constant. The approximate trajectories in the double stance phase are thus given by

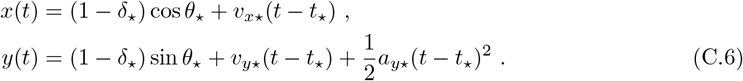

where *v*_*x*⋆_, *v*_*y*⋆_, and *a*_*y*⋆_ can be calculated at the transition time as follows:

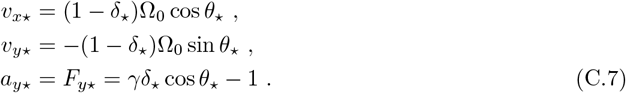

The ultimate test of these approximations, of course, will obviously be provided by comparing it with results from numerical simulation.

To summarize, Eqs. (2.5, C.5, C.6, C.7) together completely specifies a walking trajectory as a function of *λ*, Ω_0_, *γ*, and *δ*_0_. We are now going to see how to maintain a steady limit cycle gait these four parameters must obey a specific relationship that can be derived by looking at the synchronization of the periodic angular and radial motion. We will also see how different gait patterns emerge.

### D Limit cycles

#### D.1 Constraint from periodicity and synchronization

A key requirement of a sustainable walking gait is that after a given step the CoM returns to the same vertical height as the beginning of the cycle and also has the same velocity. Technically, the gait cycle should be a limit cycle. This is a technical way of ensuring that the different types of motion an animal undergoes are periodic and synchronized. For instance, in the context of the CoM motion, the vertical and horizontal motion have to be synchronized and this imposes important relationships between the parameters governing the dynamics, as we shall now see.

We will be able to derive this relationship by imposing that the time to reach the appropriate vertical and horizontal mid-step configuration that can be computed separately from the vertical and horizontal motion respectively, must be the same. For a limit cycle Using (C.6) we can calculate half of the horizontal distance traveled during the double stance:

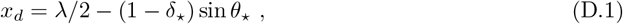

So, the half-time of the double stance phase is

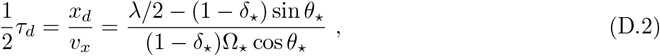

Now, due to the symmetry assumption, the vertical velocity is zero in the middle of the double stance phase. Therefore, it is possible to calculate *t*_*d*_ from the vertical kinematics as well

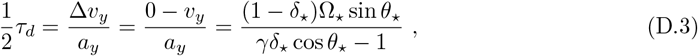

Therefore, from (D.2) and (D.3) we can conclude

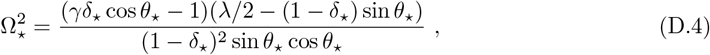

Since we suppose that the angular velocity is approximately constant during the single stance phase, we can rewrite it as

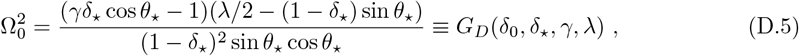

Moreover, from (2.9) we have

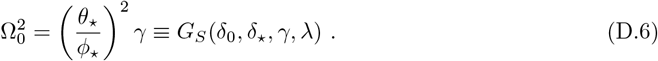

Thus, in order to have a synchronized limit cycle the four parameters, *δ*_0_, *δ*_⋆_, *γ, λ* must be related:

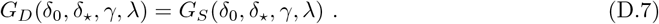

This explicitly demonstrates why all limit cycles can be characterized by only three parameters, for instance by *δ*_0_, *γ, λ*, as Ω_0_ and *δ*_⋆_ can be obtained via (D.6) and (D.7).

#### D.2 Different oscillatory mode solutions arise from the single stance phase constraint

In this subsection we will see how the gait parameter space of periodic (limit cycle) walking separates into different regions with different characteristic features. The different gaits fundamentally arise because *δ*(*t*) is a periodic function. Technically, one can see its effect in the multivalued nature of *ϕ*_⋆_ as a function of *γ, δ*_0_ and *δ*_⋆_ as inferred from (C.4) using the cosine inverse. This in turn makes *G*_*S*_ a multivalued function and choosing different branches while solving (D.7) leads to different oscillatory limit cycle gaits. To understand this more intuitively suppose one wants to travel at a given speed (approximately fixing Ω_0_) and a given step-length (*λ*). What the oscillatory evolution of *δ*(*t*) suggests is that even if one fixes the mid-stance contraction (*δ*_0_), there may be more than one way to achieve synchronization needed for limit cycle walking. For instance consider the single stance synchronization condition (D.6): One can maintain approximately the same Ω_0_, with the same transition angle ^1^, *θ*_⋆_, either by choosing a relatively lower value of *γ* and oscillating less (smaller *ϕ*_⋆_), or have a much higher *γ* and oscillate more (*ϕ*_⋆_ approximately larger by a multiple of 2*π*). To ensure that the upward velocity can be reversed during the double stance phase, the trajectory with the smaller *γ* does require a little longer double stance time as compared to the larger *γ* trajectory. So, the transition must occur a little earlier in the lower oscillatory mode, and accordingly *t*_⋆_, *δ*_⋆_, and *θ*_⋆_, are not exactly the same for the two trajectories. However, the flexibility of undergoing different phases of oscillation approximately separated by multiples of 2*π* explains how the gait parameter space separates into different oscillatory gaits, and why even with the same *λ*, Ω_0_ and *δ*_0_, different *γ* and correspondingly different oscillatory modes are possible.

### E Approximate speed range for different oscillatory gaits

In this section, we provide a technical discussion on why the different oscillatory gaits are associated with different speed ranges. We specifically demonstrate why it is not possible to walk too fast in the normal walking gait.

#### E.1 Inverted and grounded running can lead to high walking-speeds

We will first discuss the inverted walking gait whose CoM trajectory resembles that of the normal walking gait but has a different vGRF profile. We will show that while it is subjected to a lower bound in speed, one can theoretically walk much faster using this gait as compared to the normal walking gait. To see this, let us remind ourselves that for inverted walking approximately we have, 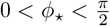.According to (D.6), for a fixed *γ* one can decrease the speed by increasing *ϕ*_⋆_, but since the latter has an upperbound leading we have

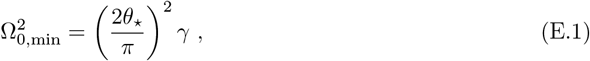

where approximately *θ*_⋆_ should be calculated by substituting *δ*_⋆_ = *δ*_*eq*_ = 1/*γ* consistent with *ϕ*_⋆_ = *π*/2. Incidentally, this coincides with the upperbound for normal walking, see also Fig.?. In contrast to having a lower bound in speed for a fixed *γ*, by decreasing *ϕ*_⋆_ all the way to zero, the speed can be increased arbitrarily according to the single stance constraint (D.6). Just as in the normal walking gait though, the velocity redirection constraint coming from the double stance phase limits the maximum speed attainable and this bound agrees well with our numerical simulation. Nevertheless, *ϕ*_⋆_ can be much smaller in the inverted walking gait in comparison with the range available for normal walking gait, and therefore much larger speeds can be accessed in this gait as compared to the normal walking gait.

Let us next focus on the grounded running gait. In contrast to all other gaits the grounded running gait has an inverted CoM trajectory where in between the mid-stance and mid-step during the single stance phase, the CoM has a vertically upward velocity. This obviates the need to have an upward force during the double stance phase in order to redirect the velocity. This means that we should no longer require *δ*_⋆_ >*δ*_*eq*_ = 1/*γ*. So, *ϕ*_⋆_ need not satisfy, 0 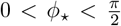,but could be larger, as borne out bt our simulations. More importantly, it is clear that in the grounded running gait, the radial velocity can no longer be ignored as compared to the angular velocity, in fact, the upward component of the radial velocity dominates over the downward component associated with the angular motion. Thus our estimate of the transition velocity (C.7), which was essentially based on angular motion, can no longer be trusted, and the limit cycle constraint (D.7) which gave rise to the maximum speed-bound in other gaits, is no longer valid. Surprisingly though our analytical estimates for such gaits continue to be broadly consistent with the numerical simulations, see Fig.?. Intuitively, high speeds in normal walking gait became impossible to attain because the upward force had a maximum and the time it had in the double stance phase shrunk with increasing speed eventually making it impossible to redirect the vertical velocity. Grounded running is this very special gait where the velocity in the single stance phase after the mid-step is upward and hence there is no need for velocity redirection. Thus the speed maximum constraint coming from velocity redirection is not applicable, and indeed in our numerical simulations we see the grounded running gait to be able to access larger and larger speeds by increasing *γ*.

#### E.2 Normal walking is bounded by the double stance phase constraint

For normal walking we have shown that 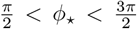. Moreover, we know that by varying *δ*_⋆_, *θ*_⋆_ does not change too much (see Figure 4B). So according to (D.6), again we have two options to increase the speed. Decreasing *ϕ*_⋆_ and increasing *γ*. However, in contrast to grounded running, there is a conflict between these two options for normal walking. In summary, for high speeds, if *ϕ*_⋆_ decreases as much as possible, we have 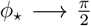, that leads to *γδ*_⋆_ → 1; so the force might not be enough to redirect the CoM velocity during double stance phase. In other words, the increase in speed needs an increase in transition force; and to have the maximum transition force we must have *γδ*_⋆_ → 2, which leads to *ϕ*_⋆_ → *π*. So at the upper bound of speed, to satisfy both constraints from single and double support phases ((D.5) and (D.6)), *ϕ*_⋆_ settles somewhere between 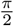 and *π*.

On the other hand, there is only a little effect of the double stance constraint on the lower bound of speed (see Figure 5G and S4). This boundary deviation from the single stance constraint can be observed better for high values of *γ* in which the need for higher force increases. For the lower bound, although *ϕ*_⋆_ is somewhere between 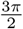 and *π*, it is much closer to 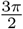 rather than *π*.

#### E.3 Slow walking via multiple oscillation modes

According to (D.6), by increasing *ϕ*_⋆_ over the normal walking range, it is quite possible to jump to the slow walking region. In this situation, since there is no concern about the speed-force relationship, the double stance constraint does not play the main role again.

### F Lower bound on stiffness and velocity constraints on gaits with multiple oscillations

#### F.1 Lowerbound on *γ* from requirement of a double stance phase

There are two other features of the gaitspace. First, requiring a finite single stance phase places a floor on *γ*. To change the vertical component of the velocity during the double stance phase at the transition

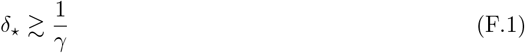

We can now use the transition geometry to find a lower bound on *γ*. As *δ*_⋆_ increases the transition occurs at smaller *θ*_⋆_ angles (2.10), also see Figure 4A. Thus if *δ*_⋆_ is pushed to a very large value by decreasing *γ, θ*_⋆_ will become zero, and there won’t be any single stance phase at all. By setting *θ*_⋆_ = 0 depicting the extreme configuration when the transition to double stance occurs at the mid-stance, we get

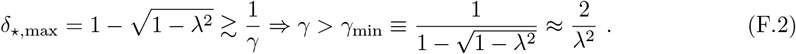

#### F.2 The analysis above extends to limit cycles with multiple oscillations

The general form of solutions of (C.4) that have a leg-length minimum (or vGRF maximum) at mid-stance can be written as

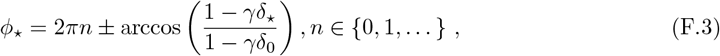

where we have assumed, *γδ*_⋆_ >1, *γδ*_0_ >1 and arccos(*ϕ*) is defined as cos^−1^ *ϕ* with *ϕ* restricted to the first quadrant, 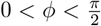. The lowest *n* = 0 mode leads to 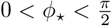 and corresponds to the inverted gaits ^2^, the most commonly observed gait among these grounded-running *like* oscillatory modes. In contrast, the normal walking gait, which exhibits a leg-length maximum (or vGRF minimum) at mid-stance is the lowest oscillatory mode (n=1) among the normal-walking *like* oscillatory gaits:

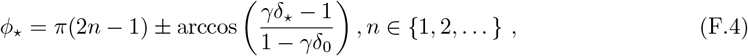

where now we have *γδ*_⋆_ >1, but *γδ*_0_ < 1. The normal walking gait can thus represent solutions with *ϕ*_⋆_ either in the second 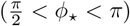 or the third 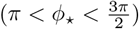 quarter of the unit circle.

The multiple branches of *ϕ*_⋆_(*δ*_⋆_) lead to multiple branches of *G*_*S*_ as a function of *δ*_⋆_, and eventually many intersections of *G*_*S*_ with *G*_*D*_. Thus we can have many limit cycles with the same speed and *δ*_0_ that, nevertheless, belong to different oscillatory gaits. Since the higher oscillatory modes correspond to lower *G*_*S*_ curves, the allowed speed range keeps decreasing as the number of oscillations increases.

There is one gait, the grounded running gait, for which the above approximate strategy fails (and is also unnecessary), as discussed in Appendix E. Essentially, in the grounded running gait, there is no longer any need to redirect the velocity in the double stance phase, and hence our analytical calculations are not valid.

**Figure S1.**
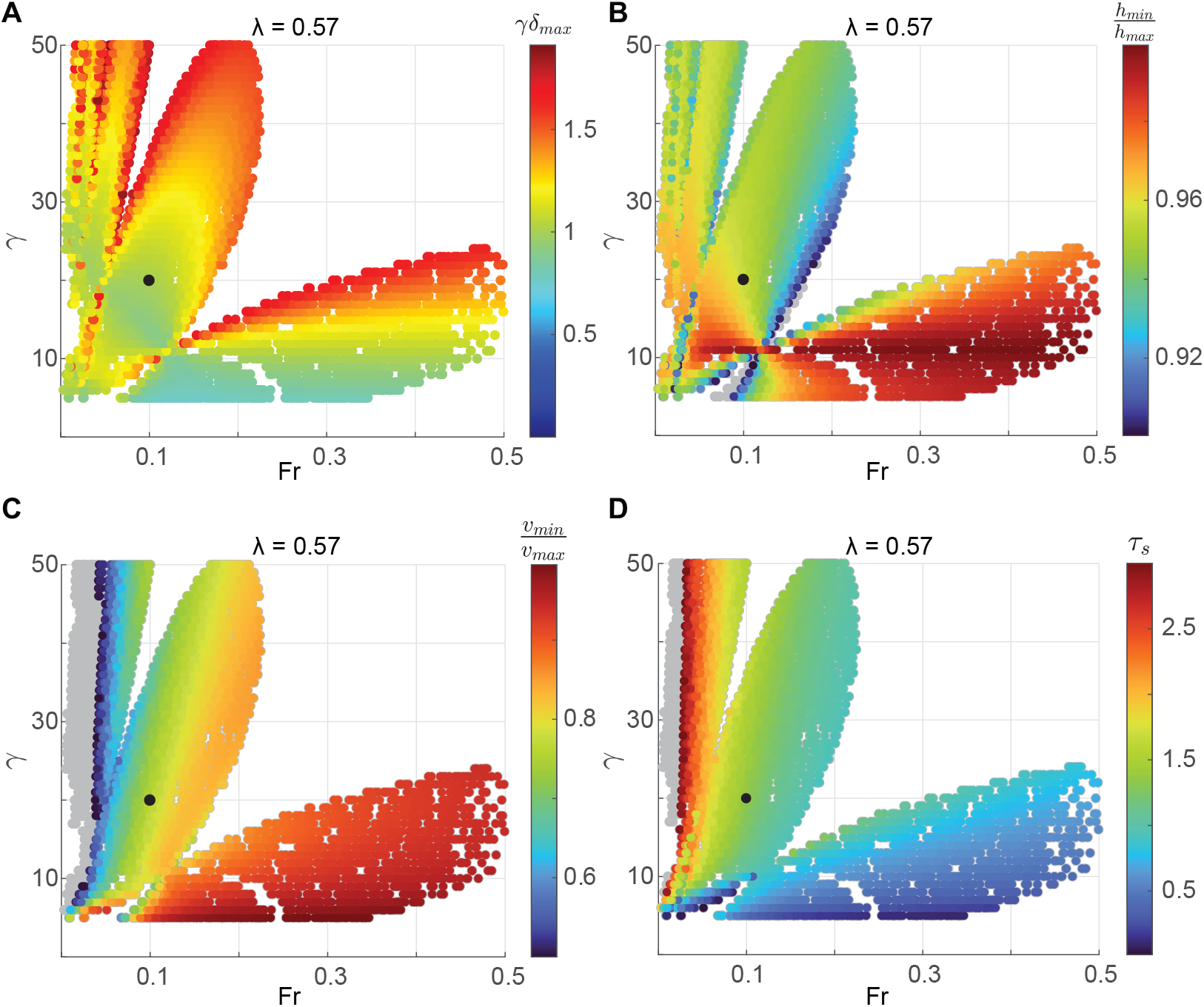
How important dynamic and kinematical features vary across gaits. **A**. We show how the maximal force, *γδ*_max_, exterted during a gait cycle varies across limit cycles. We note that lower the number of oscillations the lower is the maximal force required.In **B**. and **C**. We assess how the height and horizontal speed varies during a gait cycle by calculating the ratio between their maximum and minimum values, *h*_min_/*h*_max_, and *v*_min_/*v*_max_ respectively. We note that while the variations in the normal gait lies mostly within the ranges observed in humans, the higher oscillatory gaits show a larger variation in speed. **D**. Here we depict how the single-stance or swing time varies across different gait cycles. We see that cycles with more number of oscillations have a longer time and therefore lower frequency. Since energy loss due to swing increases with higher frequency, this suggests that high oscillatory modes are energetically preferred. In all these figures the black dot represents the limit cycle that best fits experimental walking data at 2.5 miles/hour. We note that it exhibits relatively small variation in speed and height. Moreover, as compared to inverted gait cycles (at the same speed) it expends less swing energy, and as compared to higher oscillatory modes exerts less force. In concert, these plots argue why the normal gait is the preferred gait.

**Figure S2.**
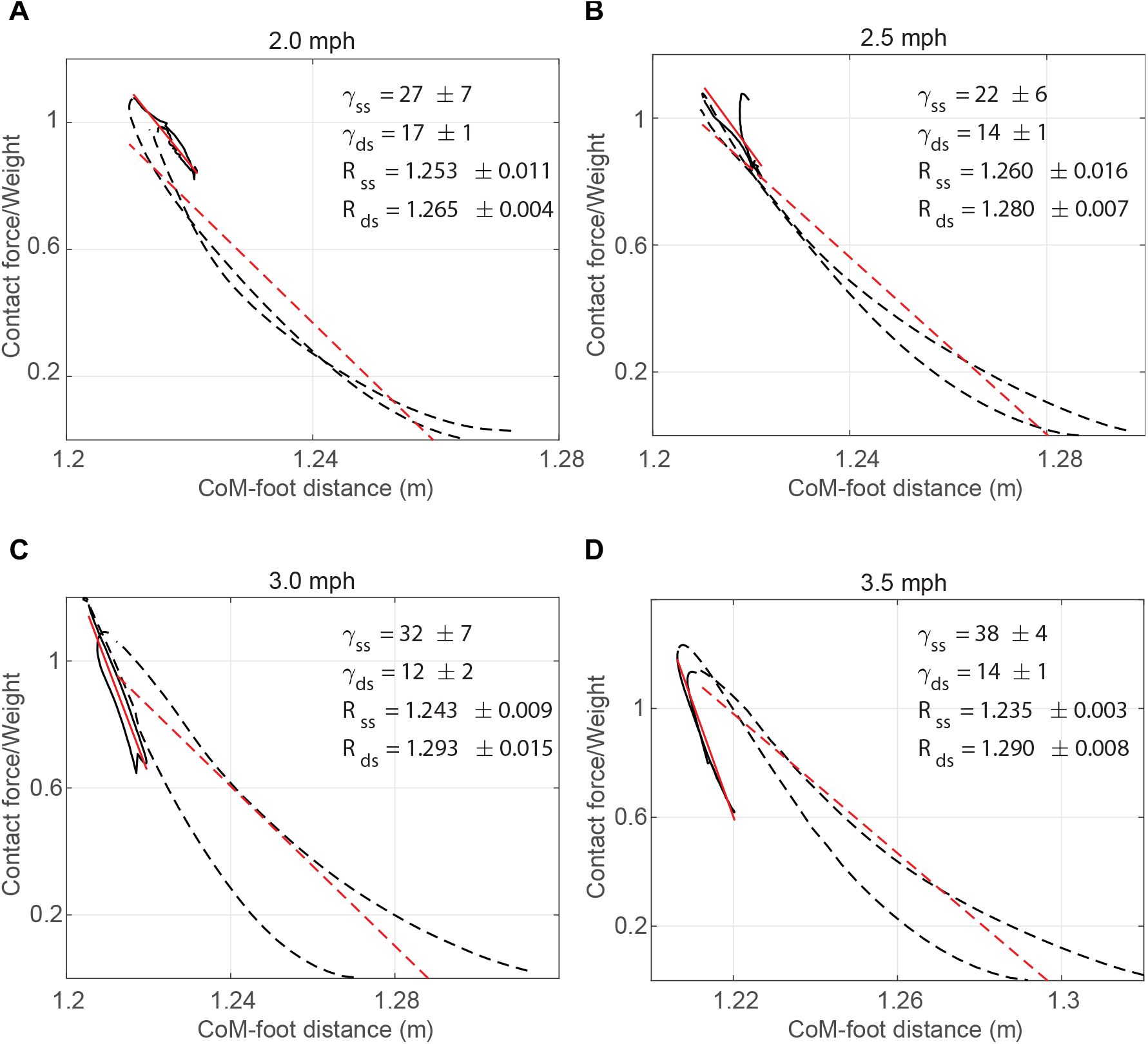
Force-length relationship shows that except for walking at 2.5 mph, the spring constants during single and double stance phases are different. Each panel shows the force-length relationship for a single step. Solid black line is during the single support phase, and dotted black lines are during the double support phase. Red solid and dotted lines show the best fitting linear spring to the single and double support phases. The mean and the SD of the spring constants and natural leg length are also reported for each speed.

**Figure S3.**
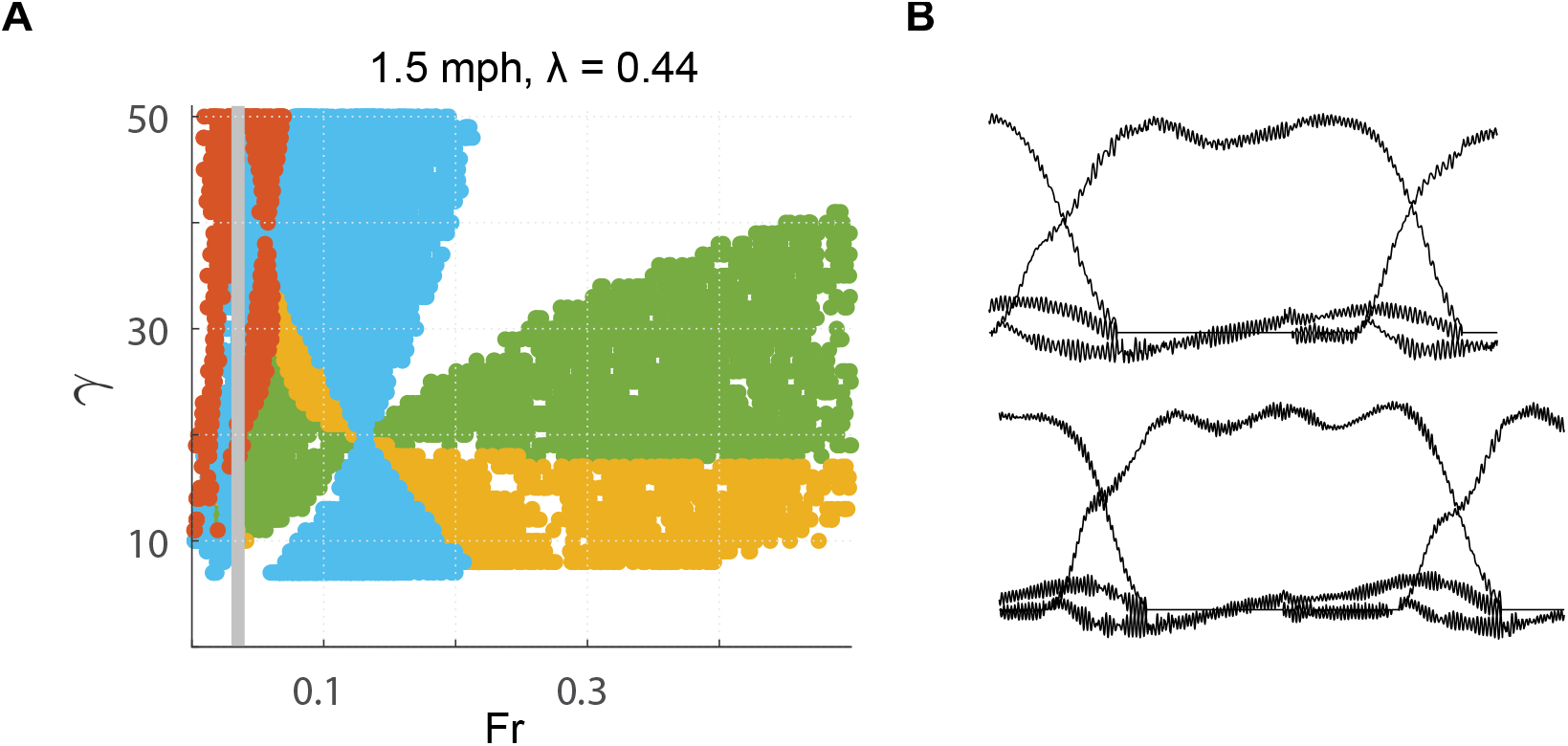
Human walking can involve higher oscillatory modes at low speed. **A**. At low speeds, such as, at a Fr number of 0.04 (gray line), both an M-shaped GRF (blue), and higher oscillation mode (orange) are possible. **B**. vGRF at these walking speeds can show both an M-shaped GRF, and GRF with higher number of oscillation as seen by the three-humped vGRF pattern.

**Figure S4.**
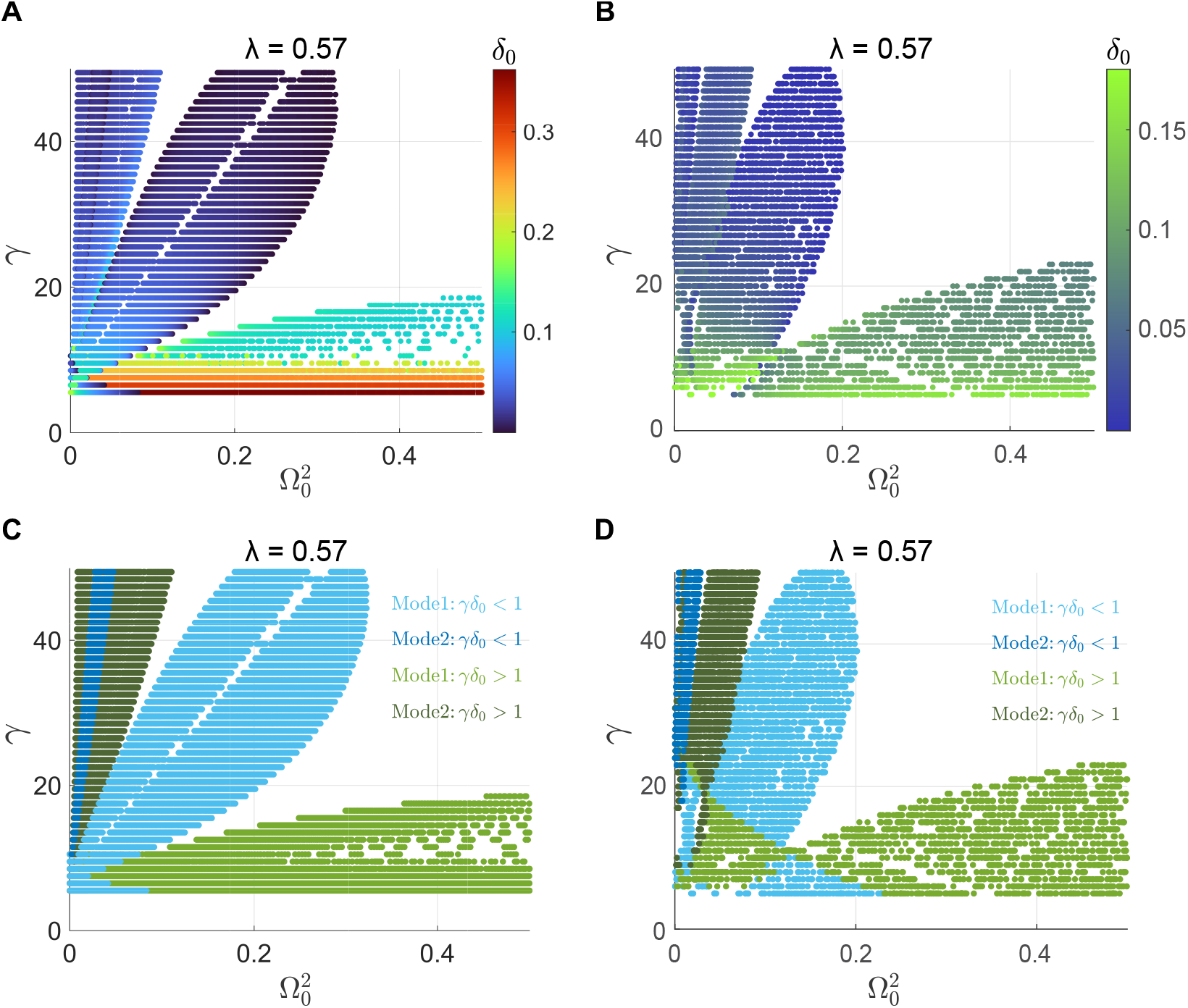
Comparison between analytically and numerically obtained limit cycles. **A**. We show the analytical solutions for a fixed step-length that are characterized by three quantities: the x-axis and y-axis corresponds to 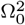 and *γ* respectively, while the color represents the value of *δ*_0_. **B**. To compare with the analytical results we here depict numerical limit cycle solutions using the same color axis scale to represent *δ*_0_ values. The analytical and numerical plots show similar patterns, and while the analytical solution over-estimates the value of *δ*_0_, its variation both along the Ω^2^-axis and 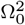-axis show similar trend as the numerical plot. **C**. and **D**. show the same plots as **A**. and **B**. respectively, except that the color now represents the identity of the gait, normal, inverted or exhibiting multiple oscillations. While there are some discrepancies between the analytical and numerical results, they are broadly consistent with each other.

## Footnotes

1 In other words, achieve approximately the same contraction length, *δ*_⋆_, approximately at the same same time, *t*_⋆_.

2 Since *ϕ*_⋆_ can only be positive, the *ϕ*_⋆_ ∈ (−*π*/2, 0) is unphysical and absent from the *n* = 0 grounded running gait.

## References

Adamczyk, P. G. and Kuo, A. D. (2009). Redirection of center-of-mass velocity during the step-to-step transition of human walking. Journal of Experimental Biology, 212(16):2668–2678.

Ahn, A. N., Furrow, E., and Biewener, A. A. (2004). Walking and running in the red-legged running frog, kassina maculata. Journal of Experimental Biology, 207(3):399–410.

Andrada, E., Blickhan, R., Ogihara, N., and Rode, C. (2020). Low leg compliance permits grounded running at speeds where the inverted pendulum model gets airborne. Journal of Theoretical Biology, 494:110227.

Andrada, E., Nyakatura, J. A., Bergmann, F., and Blickhan, R. (2013a). Adjustments of global and local hindlimb properties during terrestrial locomotion of the common quail (coturnix coturnix). Journal of Experimental Biology, 216(20):3906–3916.

Andrada, E., Rode, C., and Blickhan, R. (2013b). Grounded running in quails: simulations indicate benefits of observed fixed aperture angle between legs before touch-down. Journal of theoretical biology, 335:97–107.

Andrada, E., Rode, C., Sutedja, Y., Nyakatura, J. A., and Blickhan, R. (2014). Trunk orientation causes asymmetries in leg function in small bird terrestrial locomotion. Proceedings of the Royal Society B: Biological Sciences, 281(1797):20141405.

Antoniak, G., Biswas, T., Cortes, N., Sikdar, S., Chun, C., and Bhandawat, V. (2019). Spring-loaded inverted pendulum goes through two contraction-extension cycles during the single-support phase of walking. Biology open, 8(6):bio043695.

Basu, C., Wilson, A. M., and Hutchinson, J. R. (2019). The locomotor kinematics and ground reaction forces of walking giraffes. Journal of Experimental Biology, 222(2):jeb159277.

Biswas, T., Rao, S., and Bhandawat, V. (2018). A simple extension of inverted pendulum template to explain features of slow walking. Journal of theoretical biology, 457:112–123.

Blickhan, R. (1989). The spring-mass model for running and hopping. Journal of biomechanics, 22(11-12):1217–1227.

Blickhan, R., Andrada, E., Hirasaki, E., and Ogihara, N. (2018). Global dynamics of bipedal macaques during grounded and aerial running. Journal of Experimental Biology, 221(24):jeb178897.

Blickhan, R. and Full, R. (1993). Similarity in multilegged locomotion: bouncing like a monopode. Journal of Comparative Physiology A, 173:509–517.

Bonnaerens, S., Fiers, P., Galle, S., Aerts, P., Frederick, E. C., Kaneko, Y., Derave, W., et al. (2019). Grounded running reduces musculoskeletal loading. Medicine and Science in Sports and Exercise, 51(4):708–715.

Buczek, F. L., Cooney, K. M., Walker, M. R., Rainbow, M. J., Concha, M. C., and Sanders, J. O. (2006). Performance of an inverted pendulum model directly applied to normal human gait. Clinical Biomechanics, 21(3):288–296.

Daley, M. A., Felix, G., and Biewener, A. A. (2007). Running stability is enhanced by a proximodistal gradient in joint neuromechanical control. Journal of Experimental Biology, 210(3):383–394.

Davis, S., Fox, A., Bonacci, J., and Davis, F. (2020). Mechanics, energetics and implementation of grounded running technique: a narrative review. BMJ Open Sport & Exercise Medicine, 6(1):e000963.

Ding, J., Moore, T. Y., and Gan, Z. (2022). A template model explains jerboa gait transitions across a broad range of speeds. Frontiers in Bioengineering and Biotechnology, 10.

Donelan, J. M., Kram, R., and Kuo, A. D. (2002). Mechanical work for step-to-step transitions is a major determinant of the metabolic cost of human walking. Journal of experimental biology, 205(23):3717–3727.

Farley, C. T. and Gonzalez, O. (1996). Leg stiffness and stride frequency in human running. Journal of biomechanics, 29(2):181–186.

Gan, Z., Yesilevskiy, Y., Zaytsev, P., and Remy, C. D. (2018). All common bipedal gaits emerge from a single passive model. Journal of The Royal Society Interface, 15(146):20180455.

Geyer, H. (2005). Simple models of legged locomotion based on compliant limb behavior. PhD thesis.

Geyer, H., Seyfarth, A., and Blickhan, R. (2006). Compliant leg behaviour explains basic dynamics of walking and running. Proceedings of the Royal Society B: Biological Sciences, 273(1603):2861–2867.

Griffin, T. M., Main, R. P., and Farley, C. T. (2004). Biomechanics of quadrupedal walking: how do four-legged animals achieve inverted pendulum-like movements? Journal of Experimental Biology, 207(20):3545–3558.

Hubel, T. Y. and Usherwood, J. R. (2015). Children and adults minimise activated muscle volume by selecting gait parameters that balance gross mechanical power and work demands. Journal of Experimental Biology, 218(18):2830–2839.

Kim, S. and Park, S. (2011). Leg stiffness increases with speed to modulate gait frequency and propulsion energy. Journal of biomechanics, 44(7):1253–1258.

Kram, R., Domingo, A., and Ferris, D. P. (1997). Effect of reduced gravity on the preferred walk-run transition speed. The Journal of experimental biology, 200(4):821–826.

Kuo, A. D. (2001). A simple model of bipedal walking predicts the preferred speed–step length relationship. J. Biomech. Eng., 123(3):264–269.

Kuo, A. D. (2002). Energetics of actively powered locomotion using the simplest walking model. J. Biomech. Eng., 124(1):113–120.

Kuo, A. D., Donelan, J. M., and Ruina, A. (2005). Energetic consequences of walking like an inverted pendulum: step-to-step transitions. Exercise and sport sciences reviews, 33(2):88–97.

Lee, C. R. and Farley, C. T. (1998). Determinants of the center of mass trajectory in human walking and running. Journal of experimental biology, 201(21):2935–2944.

Lim, H. and Park, S. (2019). A bipedal compliant walking model generates periodic gait cycles with realistic swing dynamics. Journal of Biomechanics, 91:79–84.

Lin, B., Živanović, S., Zhang, S., Zhang, Q., and Fan, F. (2023). Evaluation of compliant walking locomotion models for civil engineering applications. Journal of Sound and Vibration, 561:117815.

Lipfert, S. W., Günther, M., Renjewski, D., Grimmer, S., and Seyfarth, A. (2012). A model-experiment comparison of system dynamics for human walking and running. Journal of theoretical biology, 292:11–17.

Mauersberger, M., Hähnel, F., Wolf, K., Markmiller, J. F., Knorr, A., Krumm, D., and Odenwald, S. (2022). Predicting ground reaction forces of human gait using a simple bipedal spring-mass model. Royal Society Open Science, 9(7):211582.

Maus, H.-M., Lipfert, S., Gross, M., Rummel, J., and Seyfarth, A. (2010). Upright human gait did not provide a major mechanical challenge for our ancestors. Nature communications, 1(1):70.

Maus, H.-M., Revzen, S., Guckenheimer, J., Ludwig, C., Reger, J., and Seyfarth, A. (2015). Constructing predictive models of human running. Journal of The Royal Society Interface, 12(103):20140899.

McMahon, T. A. and Cheng, G. C. (1990). The mechanics of running: how does stiffness couple with speed? Journal of biomechanics, 23:65–78.

Müller, R., Rode, C., Aminiaghdam, S., Vielemeyer, J., and Blickhan, R. (2017). Force direction patterns promote whole body stability even in hip-flexed walking, but not upper body stability in human upright walking. Proceedings of the Royal Society A: Mathematical, Physical and Engineering Sciences, 473(2207):20170404.

Nishikawa, K., Biewener, A. A., Aerts, P., Ahn, A. N., Chiel, H. J., Daley, M. A., Daniel, T. L., Full, R. J., Hale, M. E., Hedrick, T. L., et al. (2007). Neuromechanics: an integrative approach for understanding motor control. Integrative and comparative biology, 47(1):16–54.

Reinhardt, L. and Blickhan, R. (2014). Level locomotion in wood ants: evidence for grounded running. Journal of Experimental Biology, 217(13):2358–2370.

Rummel, J., Blum, Y., and Seyfarth, A. (2010). Robust and efficient walking with spring-like legs. Bioinspiration & biomimetics, 5(4):046004.

Schmitt, D. (1999). Compliant walking in primates. Journal of Zoology, 248(2):149–160.

Shorten, M. and Pisciotta, E. (2017). Running biomechanics: what did we miss? ISBS Proceedings Archive, 35(1):293.

Srinivasan, M. (2011). Fifteen observations on the structure of energy-minimizing gaits in many simple biped models. Journal of The Royal Society Interface, 8(54):74–98.

Srinivasan, M. and Ruina, A. (2006). Computer optimization of a minimal biped model discovers walking and running. Nature, 439(7072):72–75.

Usherwood, J. R. (2005). Why not walk faster? Biology letters, 1(3):338–341.

Whittington, B. R. and Thelen, D. G. (2009). A simple mass-spring model with roller feet can induce the ground reactions observed in human walking. Journal of Biomechanical Engineering, 131(1).

Wisse, M., Schwab, A. L., and van der Helm, F. C. (2004). Passive dynamic walking model with upper body. Robotica, 22(6):681–688.

